# Dual targeting of EZH2 and Histone Deacetylases in hematological malignancies promotes transcriptional and metabolic deregulation leading to ferroptosis

**DOI:** 10.1101/2024.03.03.583195

**Authors:** Alexandra Veloso, Thomas Slegers, Jente Schoenaker, Sofie Demeyer, Stef Van den Bergh, Margo Aertgeerts, Krista Verhoeft, Yilke Schoenmaekers, Nienke Meeuws, Ilan Bischops, Félicien Renard, Lien Boogaerts, Nicole Mentens, Olga Gielen, Kris Jacobs, Heidi Segers, Jan Cools, Daan Dierickx, Marlies Vanden Bempt

## Abstract

The methyltransferase EZH2 functions as the enzymatic component of the PRC2 complex, which deposits methyl groups on H3K27, leading to chromatin condensation and gene repression. Recent studies have shown that EZH2 can also act as a transcriptional modulator outside of the PRC2 complex and thus, independent of its methyltransferase activity. In this study, we first aimed to investigate the effects of EZH2 enzymatic inhibition versus protein degradation in hematological malignancies. We demonstrate that EZH2 degradation is more effective in blocking cellular proliferation compared to EZH2 enzymatic inhibition, and that EZH2 targeting upregulates the cholesterol biosynthesis pathway. Secondly, combined targeting of EZH2 and HDACs showed synergistic effects in a broader spectrum of hematological malignancies. Mechanistically, combined targeting of EZH2 and HDACs induced increased levels of H3K27 acetylation and strong upregulation of cholesterol biosynthesis. This leads to metabolic stress due to acetyl-CoA depletion, ultimately inducing ferroptotic cell death.

**Statement of significance:** We show that combined EZH2 and HDAC targeting is a promising therapeutic strategy for a broad spectrum of hematological malignancies. We uncover that EZH2 targeting induces upregulation of cholesterol biosynthesis, which is crucial for the synergistic effect with HDAC inhibition, ultimately leading to ferroptosis induction.

## Introduction

Enhancer of Zeste Homolog 2 (EZH2) is typically known as the enzymatic component of the Polycomb repressive 2 complex (PRC2), which is responsible for adding methyl groups to Histone 3 Lysine 27 (H3K27), resulting in gene repression. This is known as the canonical function of EZH2. Besides its role in the PRC2 complex, EZH2 has been described to have diverse non-canonical functions that play a role in the development of multiple cancer types. Recently, EZH2 was described to act as a transcriptional activator independent of the PRC2 complex in multiple hematological malignancies, including peripheral T cell lymphoma (PTCL), NK T cell lymphoma (NKTCL) and acute myeloid leukemia (AML)^1–4^. This non-canonical function of EZH2 was shown to be mostly independent of its methyltransferase activity, but dependent on its phosphorylation by CDK1 in PTCL^1^ or by JAK3 in NKTCL^2^.

To date, only one specific EZH2 methyltransferase inhibitor (Tazemetostat) is FDA-approved for clinical use, more specifically for the treatment of follicular lymphoma and epithelioid sarcoma. However, Tazemetostat only blocks the canonical function of EZH2, and this might not be sufficient for cancers relying on its non-canonical activity. Therefore, the recently developed EZH2 degraders^3,5,6^ could be considered in the future for the treatment of tumors with a dependency on non-canonical EZH2 activity.

Many hematological malignancies are characterized by epigenetic deregulation as a result of the abnormal activity of histone deacetylases, which can be targeted with Histone Deacetylase (HDAC) inhibitors. HDAC inhibition was shown to be effective in preclinical B-ALL models^7^, and is currently FDA-approved for the treatment of relapsed PTCL, cutaneous T cell lymphomas or multiple myeloma^8^. However, HDAC inhibitors show only a limited efficacy as monotherapy, and are associated with significant toxicities. Therefore, different strategies are currently explored to combine HDAC inhibitors with complementary drugs to enhance their overall effectiveness. Previous studies have already shown that HDAC inhibition shows synergistic effects with EZH2 enzymatic inhibition in both solid cancers, such as castration-resistant prostate cancer^9^ and non-small cell lung cancer^10^, and in certain types of hematological cancers, such as germinal-center-derived lymphomas^11^ and multiple myeloma^12^. The HDAC inhibitor Romidepsin was also found to act synergistically with an EZH2 degrader in a mouse model of MYCN-driven PTCL^1^. Based on these findings, dual targeting of EZH2 and HDACs emerges as a promising therapeutic strategy, potentially for a broad range of hematological malignancies, although the underlying mechanism remains elusive.

## Results

### Both myeloid and lymphoid malignancies are sensitive to EZH2 degradation

Recently, EZH2 has been recognized as an interesting target for the treatment of hematological malignancies. Given the recent discovery that EZH2 can both repress and activate transcription via different mechanisms, it is currently unknown which aspect of EZH2 signaling is most critical in driving cellular proliferation. We have previously shown that murine MYCN-driven PTCL is insensitive to EZH2 inhibition, but strongly sensitive to EZH2 degradation^1^. However, it remained unknown if this is applicable to a broader set of hematological malignancies. To investigate this, we selected a panel of 23 hematopoietic cell lines (**Table S1**) derived from T cell acute lymphoblastic leukemia (T-ALL), B cell acute lymphoblastic leukemia (B-ALL), acute myeloid leukemia (AML), peripheral T cell lymphoma (PTCL), cutaneous T cell lymphoma (CTCL) and diffuse large B cell lymphoma (DLBCL), on which we performed a small-scale drug screen. We used the EZH2 methyltransferase inhibitor Tazemetostat, which targets only the canonical activity of EZH2, and three recently developed EZH2 degraders (MS1943^5^, MS177^3^ and EZH2 PROTAC^6^), which will affect both canonical and non-canonical activities of EZH2. Interestingly, most cell lines seemed insensitive to EZH2 methyltransferase inhibition, even at high concentrations after 72h of treatment, indicating that they are not dependent on the canonical function of EZH2 within the PRC2 complex. In sharp contrast, most cell lines were sensitive to EZH2 degradation (**Fig. 1A, Fig. S1, S2A**), which was not correlated with EZH1 or EZH2 expression levels (**Fig. S2B**). In contrast to EZH2 enzymatic inhibition, EZH2 degradation potently induced cell death in cell lines and *ex vivo* PDX models from both myeloid and lymphoid lineage (**Fig. 1B,C, Fig. S2C**). EZH2 enzymatic inhibition only moderately affected cell viability and proliferation, even over a longer period of time. Nevertheless, H3K27me3 levels were already strongly decreased after 72 hours of treatment and this was maintained or even further decreased after 9 days of treatment (**Fig. 1D,E, Fig. S2D**). We observed only minor toxicities to healthy peripheral blood mononuclear cells (PBMCs) after EZH2 degradation or EZH2 enzymatic inhibition (**Fig. 1F, G**), except for the EZH2 degrader MS177, which showed higher levels of toxicity.

**Figure 1:**
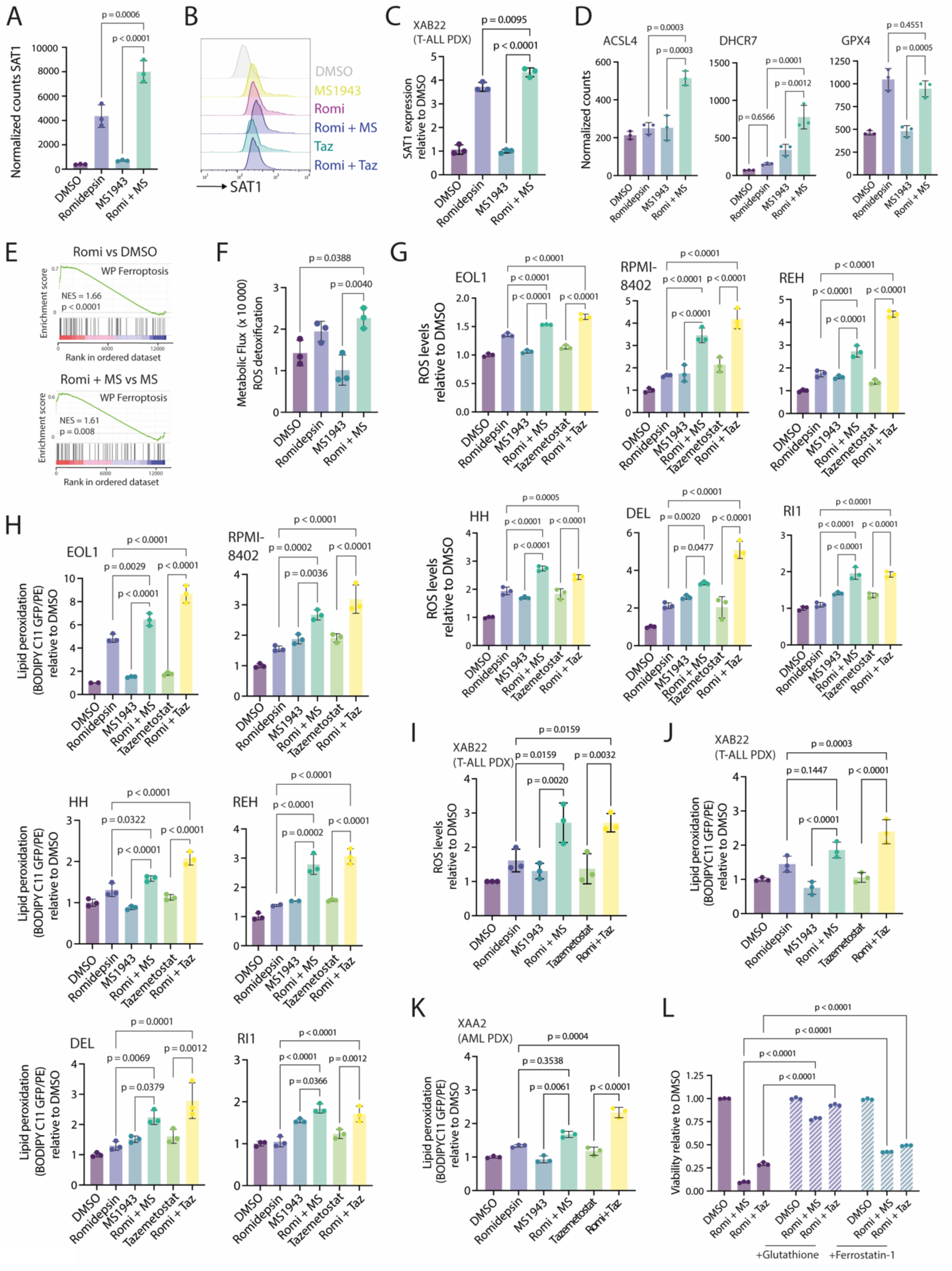
Both myeloid and lymphoid malignancies are sensitive to EZH2 degradation. A) Viability relative to DMSO of different human cell lines treated with 10 µM of EZH2 enzymatic inhibitor (Tazemetostat) or EZH2 degraders (MS177, MS1943, EZH2 PROTAC) for 48-72h. B) Annexin V / PI staining of the KG1, Jurkat, REH and RI1 cell lines after treatment for 48h with an EZH2 enzymatic inhibitor (Tazemetostat) or EZH2 degraders (MS177, MS1943). C) Annexin V staining of different PDX models after *ex vivo* treatment (24h) with an EZH2 enzymatic inhibitor (Tazemetostat, 5 µM) or an EZH2 degrader (MS1943, 5µM). D) Viability relative to DMSO of different cell lines treated for 10 days with 10 µM Tazemetostat or DMSO. E) Intracellular H3K27me3 levels after 3 (top) or 9 (bottom) days of treatment with 10 µM Tazemetostat or DMSO. E) Annexin V / PI staining of human healthy PBMCs after 24h, 48h or 96h treatment with different EZH2-targeting drugs (5 µM). F) Flow cytometric analysis of the human healthy PBMC populations treated with the EZH2 inhibitors and degraders (5 µM) for 24h or 48h.

### EZH2 targeting induces upregulation of cholesterol biosynthesis

To investigate the differential effects of EZH2 degradation versus EZH2 enzymatic inhibition on the transcriptomic landscape, we performed a transcriptomic analysis of RPMI-8402 cells treated with the EZH2 degrader MS1943 (16h) or with the EZH2 inhibitor Tazemetostat (72h). EZH2 degradation induced downregulation of previously described non-canonical EZH2 target genes (Kamminga EZH2 targets)^1,13^, and of MYC target genes, which are both gene sets associated with active proliferation. However, EZH2 degradation did not affect canonical PRC2 target genes (**Fig. 2A, B**). In contrast, enzymatic inhibition of EZH2 induced upregulation of canonical PRC2 target genes but had no effect on the non-canonical EZH2 target genes and induced a milder downregulation of MYC target genes (**Fig. 2C, D**).

**Figure 2:**
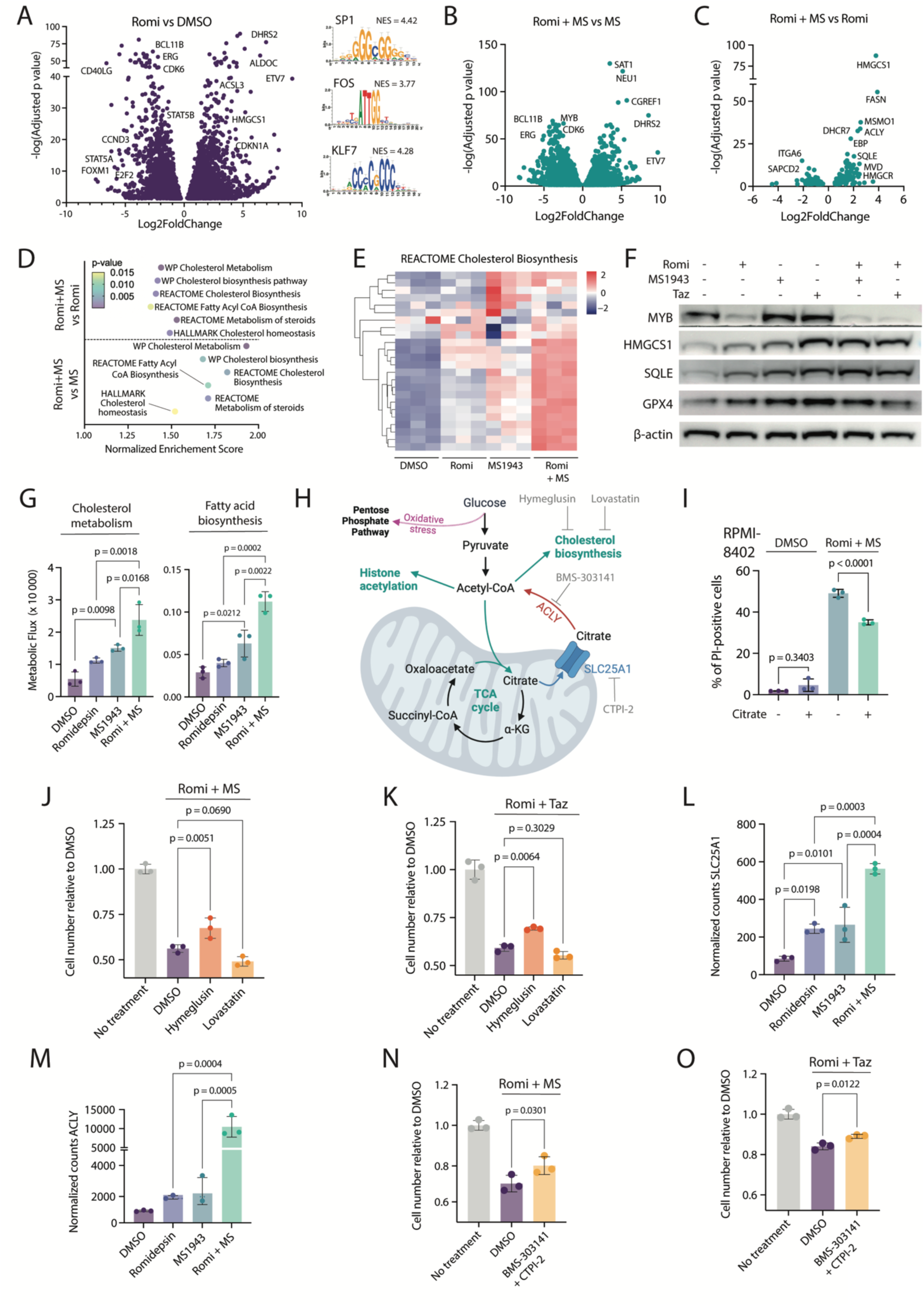
Transcriptomic analysis of EZH2 degradation and EZH2 enzymatic inhibition. A) Volcano plot showing the differentially expressed genes in RPMI-8402 cells treated with 5 µM MS1943 for 16h compared to DMSO. Enriched motifs found in the regulatory regions of the differentially expressed genes are shown on the right. B) Gene set enrichment analysis (GSEA) on the differentially expressed genes in RPMI-8402 cells treated with 5 µM MS1943 for 16h compared to DMSO. C) Volcano plot showing the differentially expressed genes in RPMI-8402 cells treated with 5 µM Tazemetostat for 72h compared to DMSO. Enriched motifs found in the regulatory regions of the differentially expressed genes are shown on the right. D) Gene set enrichment analysis (GSEA) on the differentially expressed genes in RPMI-8402 cells treated with 5 µM Tazemetostat for 72h compared to DMSO. E) Volcano plot showing the differentially expressed genes in RPMI-8402 cells treated with 5 µM MS1943 for 16h compared to treatment with 5 µM Tazemetostat for 72h. Enriched motifs found in the regulatory regions of the differentially expressed genes are shown at the bottom. F) Gene set enrichment analysis (GSEA) on the differentially expressed genes in RPMI-8402 cells treated with 5 µM MS1943 for 16h compared to treatment with 5 µM Tazemetostat for 72h. G) Heatmap with unsupervised clustering showing the expression levels of genes related to cholesterol homeostasis in RPMI-8402 cells treated with DMSO, MS1943 (5µM for 16h) or Tazemetostat (5 µM for 72h). H) qRT-PCR analysis of cholesterol biosynthesis gene expression in RPMI-8402 cells treated with 5 µM EZH2 gapmer for 48h. I) qRT-PCR analysis of cholesterol biosynthesis gene expression in ALL SIL cells treated with 4 µM EZH2 gapmer for 72h. J) qRT-PCR analysis of cholesterol biosynthesis gene expression in RPMI-8402 cells treated with 500 nM MS177 for 16h.

Overall, by comparing the differential effects of the two EZH2 targeting approaches (degradation versus enzymatic inhibition), we could show that EZH2 degradation induced stronger downregulation of MYC targets, gene sets implicated in proliferation, and non-canonical EZH2 targets, compared to EZH2 enzymatic inhibition. Remarkably, both EZH2 degradation and enzymatic inhibition induced upregulation of genes implicated in fatty acid and cholesterol biosynthesis, although this was more pronounced in the case of EZH2 degradation (**Fig. 2E-G, Fig. S2E,F**). We confirmed the effect of EZH2 loss on fatty acid and cholesterol synthesis using EZH2 gapmers (**Fig. 2H, I**) and an additional EZH2 degrader (**Fig. 2J**).

EZH2 degradation mostly affected genes characterized by EZH2 binding in regions enriched with H3K27ac, but without Suz12 and H3K27me3. However, it had little effect on canonical EZH2 target genes, characterized by co-binding of Ezh2 and Suz12 and high levels of H3K27me3. In contrast, EZH2 enzymatic inhibition induced upregulation of both canonical and non-canonical EZH2 targets (**Fig. S2G-I**).

### Dual targeting of EZH2 and HDACs induces cell death in myeloid and lymphoid malignancies

HDACs are aberrantly expressed in several hematological malignancies and have become interesting targets for clinical therapies. Indeed, HDAC inhibitors are currently used in clinical practice for the treatment of relapsed T cell lymphomas. However, their efficacy is debated, and they are associated with toxic side effects^8^. To overcome these limitations, we previously combined HDAC inhibition with EZH2 targeting, and demonstrated a synergistic effect in murine T cell lymphoma^1^. We now set out to investigate if this combination treatment is effective in a broader spectrum of acute leukemias and non-Hodgkin lymphomas.

We treated a selection of 23 cell lines derived from different hematological malignancies with the HDAC inhibitor Romidepsin combined with the EZH2 degrader MS1943 or the EZH2 inhibitor Tazemetostat. We observed a synergistic effect between EZH2 degradation and HDAC inhibition in most cell lines, independent of the immunophenotype (**Fig. 3A, Fig. S3)**. This effect was also observed using another EZH2 degrader (MS177, **Fig. S4A-D**) or other clinically approved HDAC inhibitors (Belinostat, Panobinostat) **(Fig. S4E-I**). Intriguingly, HDAC inhibition could strongly sensitize the cells for EZH2 enzymatic inhibition (**Fig. 3A**), even in case of very low single-agent sensitivity. This is in line with earlier findings in germinal-center-derived lymphomas^11^ and multiple myeloma^12^. Combined targeting of EZH2 and HDAC induced a block in the G2M phase of the cell cycle (**Fig. 3B**) and had an even stronger effect in inducing cell death compared to the single treatments in cell lines (**Fig. 3C-D**) and in *ex vivo* PDX models (**Fig. 3E**). Combined targeting of EZH2 and HDAC did not result in higher toxicities in healthy human PBMCs compared to HDAC treatment alone (**Fig. 3F, G**).

**Figure 3:**
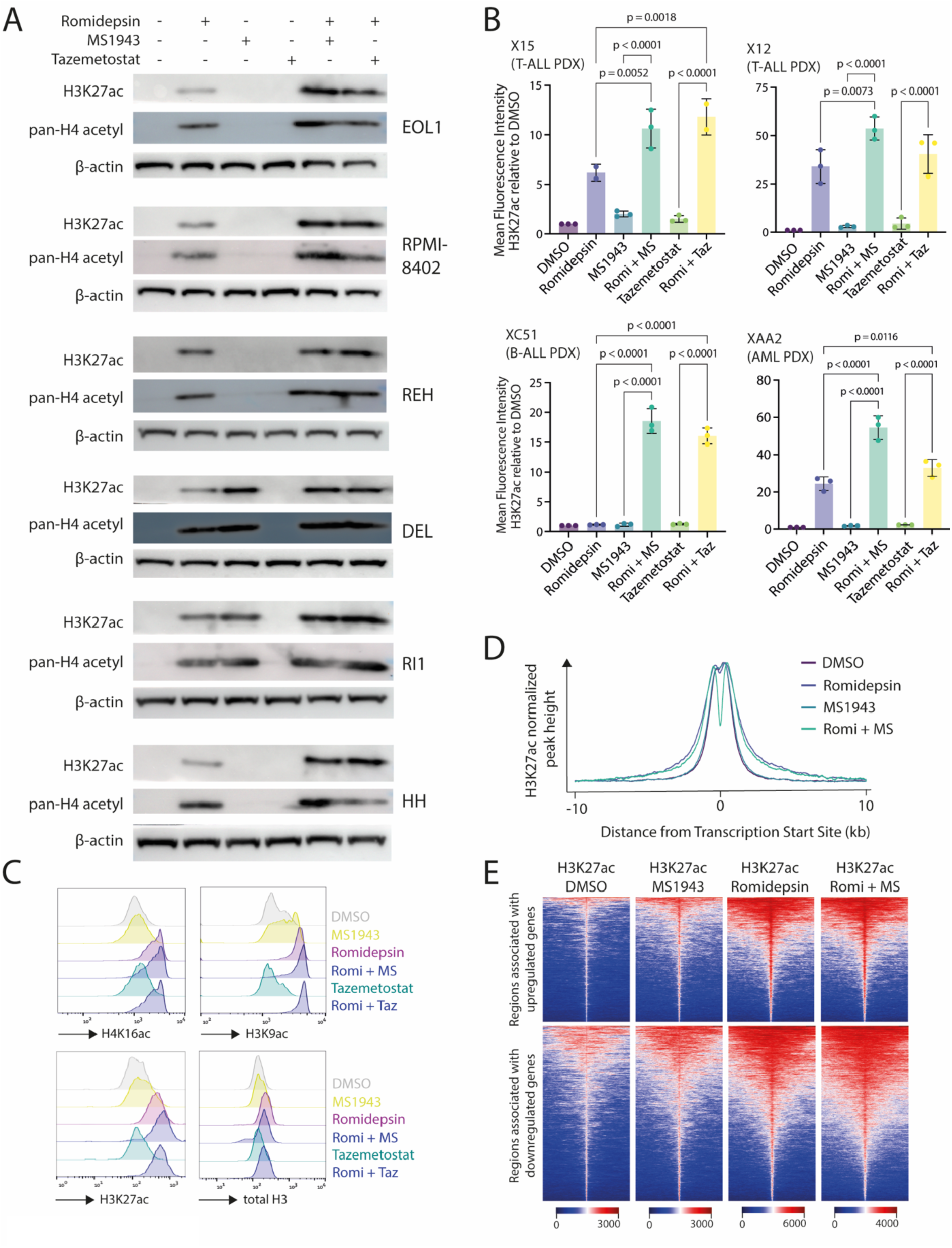
Dual targeting of EZH2 and HDACs induces cell death in myeloid and lymphoid malignancies. A) Heatmap showing the viability relative to DMSO of different human cell lines from different hematological malignancies treated with Romidepsin (3 nM), MS1943 (5 µM), Tazemetostat (5 µM) or a combination for 48h. MS = MS1943, Taz= Tazemetostat. B) Cell cycle analysis of different cell lines treated with Romidepsin (3 nM), MS1943 (5 µM), Tazemetostat (5 µM) or a combination for 48h. C) Annexin V staining of different cell lines after treatment with Romidepsin (3 nM), MS1943 (5 µM) or a combination for 24h. D) Annexin V staining of different cell lines after treatment with Romidepsin (3 nM), Tazemetostat (5 µM) or a combination for 24h. E) Annexin V staining of different PDX models after *ex vivo* treatment with Romidepsin (1-3 nM), MS1943 (5 µM), Tazemetostat (5 µM) or a combination for 24h (X15 and XB41) and 48h (XF100 and XAA2). F) Annexin V / PI staining of human healthy PBMCs after 24h, 48h or 96h treatment with Romidepsin (3 nM), MS1943 (5 µM), Tazemetostat (5 µM) or a combination. G) Flow cytometric analysis of the human healthy PBMC populations treated with the single or combination treatment for 24h or 48h.

Altogether, these data indicate that the combination of the EZH2 degrader MS1943 and the HDAC inhibitor Romidepsin lead to a stronger effect inducing leukemia cell death, yet this combination exhibits minimal toxicity towards healthy human cells, suggesting a high potential as a therapeutic treatment.

### Combined targeting of EZH2 and HDAC increases global H3K27ac levels

To investigate the molecular mechanism underlying the observed synergistic effects of combined EZH2 and HDAC targeting, we determined the global acetylation levels of Histone 3 Lysine 27 and of Histone 4 after treatment. As expected, HDAC inhibition alone already increased global histone acetylation levels. Remarkably, we observed an even stronger increase in global H3K27ac levels after combined EZH2 and HDAC targeting in cell lines (**Fig. 4A, Fig. S4J,K)** and in *ex vivo* PDX models (**Fig. 4B, Fig. S4L**), suggesting that EZH2 targeting leads to an amplification of the transcriptional deregulation caused by HDAC inhibition. In addition, HDAC inhibition also induced global increases in H4K16ac and H3K9ac, but only the latter histone mark showed a slightly more pronounced induction after combined EZH2 and HDAC targeting (**Fig. 4C**). The global increase in H3K27ac resulted in a redistribution of the H3K27ac signal with a wider spread around the transcription start site of deregulated genes (**Fig. 4D, E**) and over the gene body (**Fig. S4M**).

**Figure 4:**
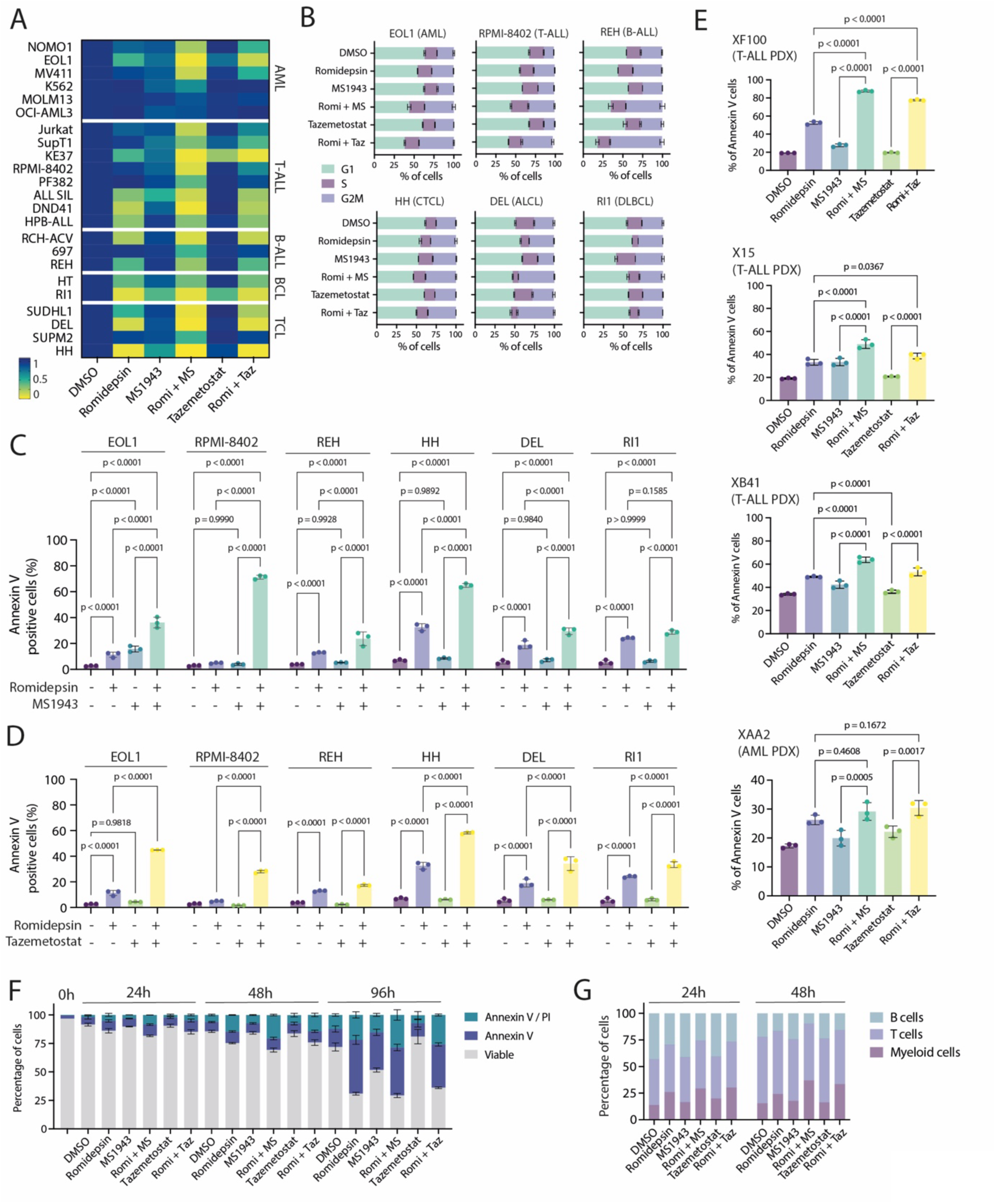
Dual targeting of EZH2 and HDACs increases global H3K27ac levels. A) Western Blot showing H3K27ac levels in different cell lines after treatment with Romidepsin (3 nM), Tazemetostat (5 µM) or a combination for 16h. B) Mean Fluorescent Intensity of the H3K27ac signal relative to DMSO in different PDX models after treatment with Romidepsin (2-5 nM), MS1943 (5 µM), Tazemetostat (5 µM) or a combination for 24h (T-ALL and B-ALL PDX models) or 48h (AML PDX model). C) Intracellular staining of H4K16ac, H3K9ac and total Histone 3 in RPMI-8402 cells treated with Romidepsin (3 nM), MS1943 (5 µM), Tazemetostat (5 µM) or a combination for 24h. D) Normalized H3K27ac signals around the transcription start site of differentially expressed genes in RPMI-8402 cells treated with Romidepsin (3 nM), MS1943 (5 µM), or a combination for 16h. E) Centered read-density heatmaps showing H3K27ac signal in RPMI-8402 cells treated with Romidepsin (3 nM), MS1943 (5 µM), or Romidepsin + MS1943 for 16h.

### Combined targeting of EZH2 and HDAC induces activation of cholesterol biosynthesis

Next, we investigated the effect of increased global H3K27ac induced by the combination treatment on the transcriptomic landscape. HDAC inhibition specifically induced downregulation of anti-apoptotic genes, such as BCL2 and BCL2L1, and of the transcription factors MYB and BCL11B (**Fig. 5A, Fig. S5A-G**). Combining EZH2 degradation with HDAC inhibition heightened the induction of transcriptional deregulation (**Fig. 5B, C**). Interestingly, we observed a more potent upregulation of genes associated with cholesterol biosynthesis and fatty acid synthesis in the combination treatment compared to the single treatments (**Fig. 5D-F, Fig. S5H, I**). Metabolic Flux balance analysis, using the computational tool METAFlux^14^, also predicted increased pathway activity of cholesterol metabolism and fatty acid biosynthesis (**Fig. 5G**) after targeting both EZH2 and HDAC (MS+ Romi).

**Figure 5:**
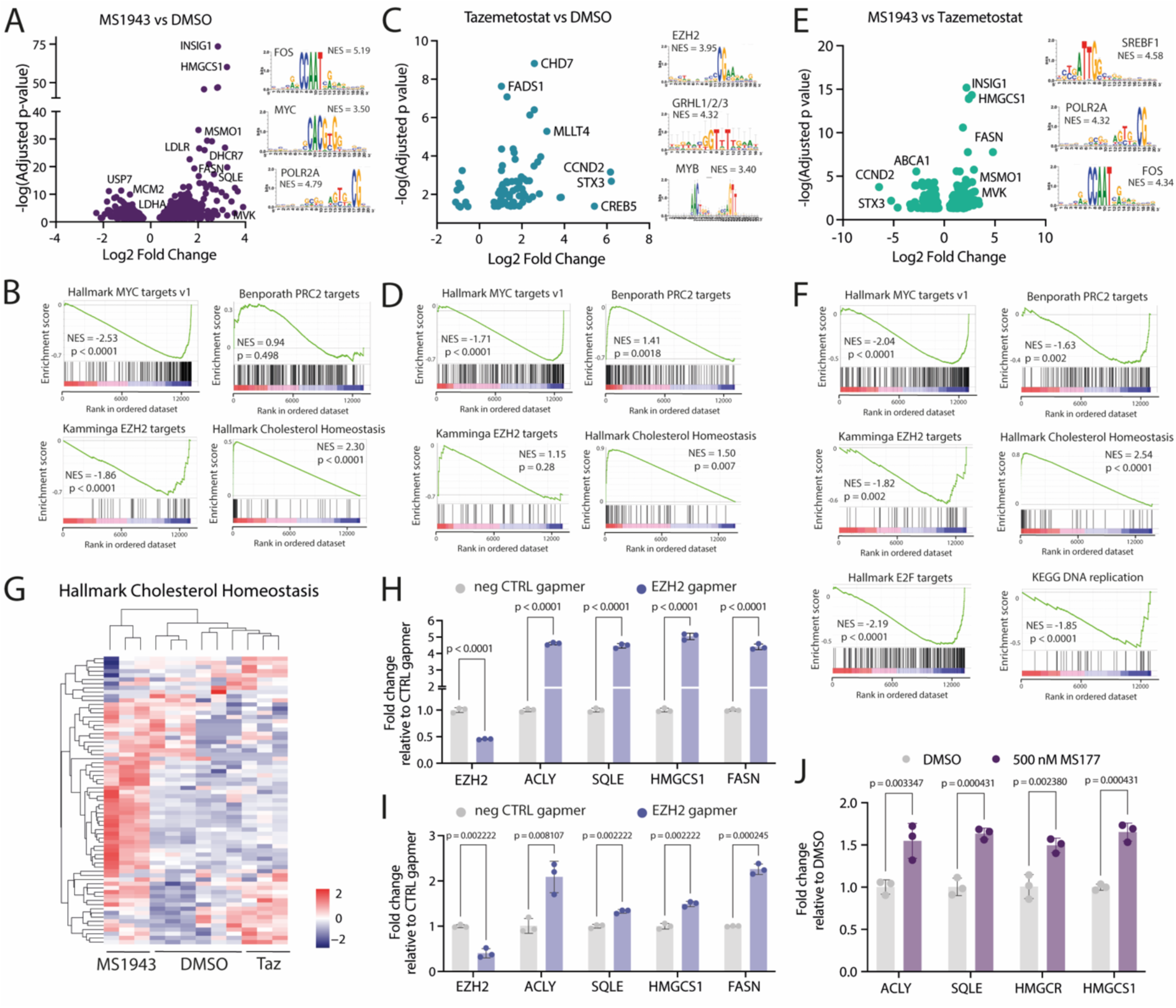
Combined targeting of EZH2 and HDAC induces activation of cholesterol biosynthesis. A) Volcano plot showing the differentially expressed genes in RPMI-8402 cells treated with Romidepsin (3 nM) for 16h compared to DMSO. Enriched motifs found in the regulatory regions of the differentially expressed genes are shown at the right. B) Volcano plot showing differentially expressed genes in RPMI-8402 cells treated with Romidepsin (3 nM) + MS1943 (5 µM) for 16h compared to treatment with MS1943 alone. C) Volcano plot showing the differentially expressed genes in RPMI-8402 cells treated with Romidepsin (3 nM) + MS1943 (5 µM) for 16h compared to treatment with Romidepsin alone. D) Normalized enrichment scores for different gene sets related to cholesterol/fatty acid metabolism in the differentially expressed genes in RPMI-8402 cells treated with DMSO, Romidepsin (3 nM), MS1943 (5 µM) or the combination for 16h. E) Heatmap showing the expression levels of the genes in the REACTOME Cholesterol Biosynthesis gene set in RPMI-8402 cells treated with DMSO, Romidepsin (3 nM), MS1943 (5µM), or the combination for 16h. F) Western Blot showing expression of MYB, HMGCS1, SQLE, and GPX4 in RPMI-8402 cells treated with DMSO, Romidepsin (3 nM), MS1943 (5 µM) or the combination for 16h. G) Pathway activation scores (∼Metabolic flux) of cholesterol metabolism and fatty acid biosynthesis as inferred using the METAFlux R package in RPMI-8402 cells treated with DMSO, Romidepsin (3 nM), MS1943 (5 µM) or the combination for 16h. H) Schematic overview of the metabolic pathways relying on acetyl-CoA. I) Percentage of PI-positive RPMI-8402 cells after 48h treatment with 3 nM Romidepsin and 5 µM MS1943, with or without 5 mM citrate supplementation. J) Relative cell number of RPMI-8402 cells treated with 3 nM Romidepsin and 5 µM MS1943 in combination with 2,5 µM Hymeglusin or Lovastatin. K) Relative cell number of RPMI-8402 cells treated with 3 nM Romidepsin and 5 µM Tazemetostat in combination with 2,5 µM Hymeglusin or Lovastatin. L) Normalized counts of *SLC25A1* in RPMI-8402 cells treated with DMSO, Romidepsin (3 nM), MS1943 (5 µM) or the combination for 16h. M) Normalized counts of *ACLY* in RPMI-8402 cells treated with DMSO, Romidepsin (3 nM), MS1943 (5 µM) or the combination for 16h. N) Relative cell number of RPMI-8402 cells treated with 3 nM Romidepsin and 5 µM MS1943 in combination with 20 µM BMS-303141 (ACLY inhibitor) + 20 µM CTPI-2 (SCL25A1 inhibitor). O) Relative cell number of RPMI-8402 cells treated with 3 nM Romidepsin and 5 µM Tazemetostat in combination with 20 µM BMS-303141 (ACLY inhibitor) + 20 µM CTPI-2 (SCL25A1 inhibitor).

As both histone acetylation and cholesterol biosynthesis solely rely on the availability of acetyl Coenzyme A (acetyl-CoA)^15^ (**Fig. 5H**), simultaneous overactivation of these processes could lead to depletion of this important metabolite from other metabolic routes, such as the tricarboxylic acid (TCA) cycle. This could result in metabolic stress, low cellular energy levels and subsequent cell death. Next, we aimed to rescue the effect of the combination treatment by supplementing the cell with the acetyl-CoA precursor citrate or by blocking the use of acetyl-CoA in cholesterol biosynthesis. First of all, citrate supplementation could indeed partially rescue the effect of the combination treatment (**Fig. 5I**). Secondly, by blocking the first step of cholesterol synthesis, the generation of HMG-CoA from acetyl-CoA and acetoacetyl-CoA, with an HMG-CoA-synthase inhibitor (Hymeglusin), we could partially rescue the toxic effects of the combination treatment. In contrast, by blocking the subsequent step in the cholesterol synthesis pathway, which consists in the reduction of HMG-CoA into mevalonate, with an inhibitor of HMGCR (Lovastatin), the toxic effect of the combination treatment was not rescued (**Fig. 5J, K**), as this does not prevent acetyl-CoA shuttling into the cholesterol synthesis pathway.

Cholesterol synthesis and histone acetylation both take place in the nucleo-cytosolic space, while the TCA cycle takes place in the mitochondria (**Fig. 5H**). Intriguingly, after combined EZH2 and HDAC targeting, we observed upregulation of the mitochondrial citrate-malate transporter SLC25A1 (**Fig. 5L**), which is responsible for shuttling acetyl-CoA-derived citrate from the mitochondria to the cytoplasm. Here, citrate is reconverted into acetyl-CoA by ATP citrate lyase (ACLY), which is also strongly upregulated by the combination treatment (**Fig. 5M**). This suggests that the combination treatment further promotes acetyl-CoA depletion from the TCA cycle by shuttling it from the mitochondria back to the cytoplasm. By blocking citrate transport to the cytoplasm through SLC25A1 (via CTPI-2 treatment) and the subsequent citrate conversion into acetyl-CoA by ACLY (via BMS-303141 treatment), we could also partially rescue the lethal effect of the combination treatment (**Fig. 5N, O**).

Altogether, these data indicate that combined EZH2 and HDAC targeting leads to lethal acetyl-CoA depletion in the cell, by shuttling it towards excessive histone acetylation and cholesterol synthesis.

### Combined targeting of EZH2 and HDAC leads to higher ROS levels and lipid peroxidation, ultimately inducing ferroptosis

High rates of fatty acid and sterol synthesis deplete the available antioxidants in the cell for reactive oxygen species (ROS) scavenging. This could lead to high intracellular ROS levels, which can in turn activate lipid peroxidation, and ultimately induce ferroptosis. In line with this, after combined EZH2 and HDAC targeting we observed upregulation of SAT1, which is a potent inducer of ferroptosis^16^ (**Fig. 6A-C**), of the ferroptosis biomarker ACSL4^17^, the pro-ferroptotic gene DHCR7^18,19^ (**Fig. 6D, Fig. S6A,B**) and of a gene set related to ferroptosis (**Fig. 6E**). In contrast, there was no additional upregulation of GPX4, the glutathione peroxidase that is the main effector of the canonical surveillance mechanism against ferroptosis^20^ after combined EZH2 and HDAC targeting (**Fig. 5F**, **Fig. 6D**). Furthermore, metabolic flux balance analysis^14^ inferred a higher activity of the ROS detoxification pathway (**Fig. 6F**), of the pentose phosphate pathway (**Fig. S6C, D**) and of glutathione metabolism (**Fig. S6E**), indicating that the combined EZH2 and HDAC targeting triggers stronger metabolic responses to oxidative stress. Altogether, these data suggest that the higher potency of combined EZH2 and HDAC targeting could be the result of a more efficient induction of oxidative stress leading to ferroptosis. Indeed, we observed significantly higher ROS levels (**Fig. 6F**) and higher levels of lipid peroxidation (**Fig. 6G**) after combined EZH2 and HDAC targeting compared to the single treatments, and we confirmed this in PDX models of both lymphoid and myeloid origin (**Fig. 6H-J, Fig. S6C**). Finally, treatment with glutathione or with the synthetic antioxidant ferrostatin-1 could rescue the toxic effects of the combination treatment (**Fig. 6K, Fig. S6D,E**). In conclusion, these results show that the synergistic effect of combined HDAC and EZH2 targeting is due to a higher level of ferroptosis induction compared to the single drugs.

**Figure 6:**
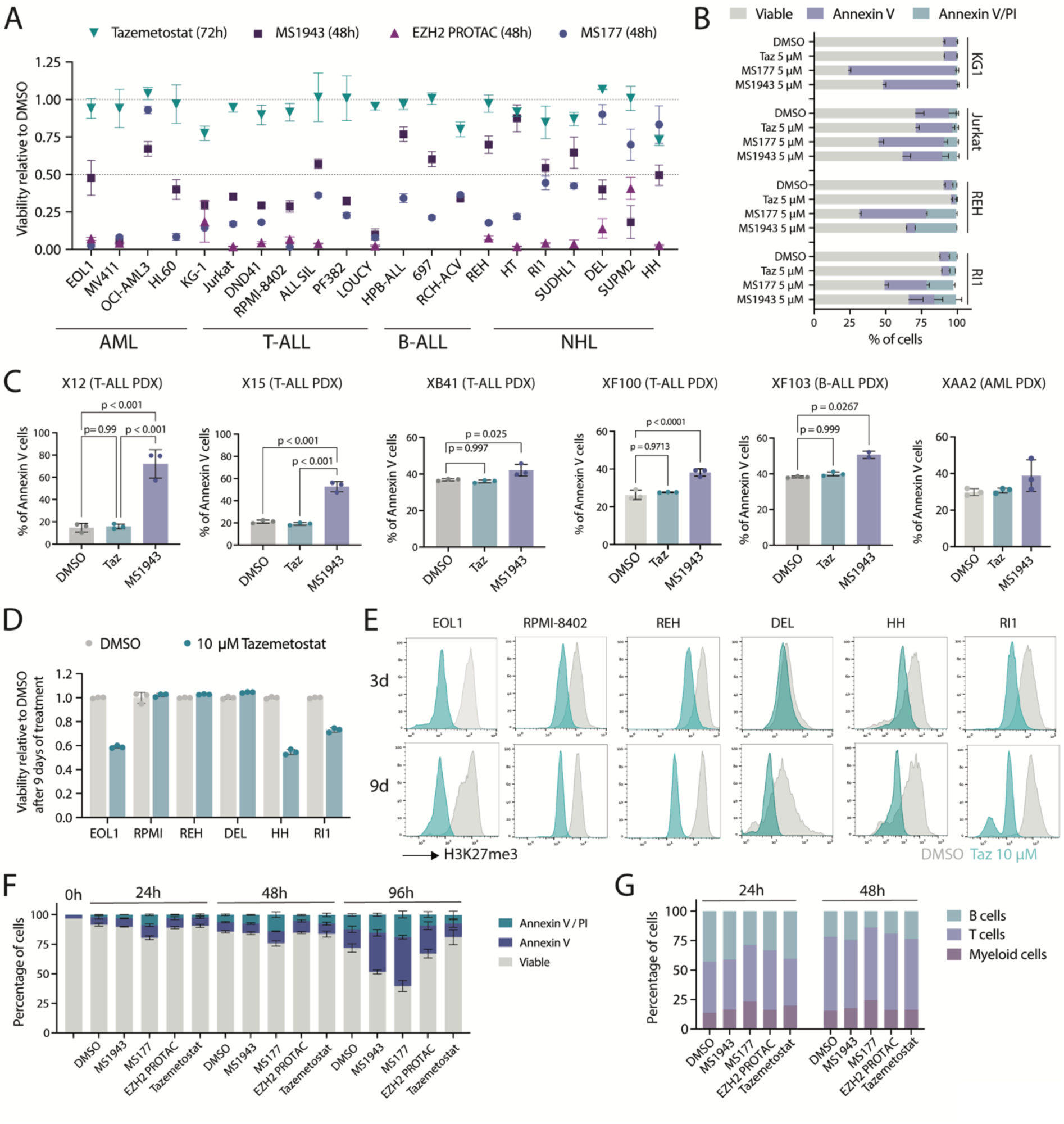
Combined targeting of EZH2 and HDAC leads to ferroptosis induction. A) Normalized counts of *SAT1* in RPMI-8402 cells treated with DMSO, Romidepsin (3 nM), MS1943 (5 µM) or the combination for 16h. B) Intracellular staining showing SAT1 levels in RPMI-8402 cells treated with DMSO, Romidepsin (3 nM), MS1943 (5 µM) or the combination for 16h. C) qRT-PCR analysis of SAT1 expression levels in XAB22 T-ALL PDX cells treated *ex vivo* with DMSO, Romidepsin (2 nM), MS1943 (5 µM) or the combination for 24h. D) Normalized counts of *ACSL4, DHCR7* and *GPX4* in RPMI-8402 cells treated with DMSO, Romidepsin (3 nM), MS1943 (5 µM) or the combination for 16h. E) Gene set enrichment analysis (GSEA) on the differentially expressed genes in RPMI-8402 cells treated with Romidepsin (3 nM) compared to DMSO (upper panel) or RPMI-8402 cells treated with Romidepsin (3 nM) in combination with MS1943 (5 µM) compared to MS1943 alone (lower panel). All treatments were performed for 16h. MS = MS1943. F) Pathway activation scores (∼Metabolic flux) of the ROS detoxification pathway as inferred using the METAFlux R package in RPMI-8402 cells treated with DMSO, Romidepsin (3 nM), MS1943 (5 µM) or the combination for 16h. G) ROS levels in different human cell lines after treatment with Romidepsin (3 nM), MS1943 (5 µM), Tazemetostat (5 µM) or a combination for 24h. H) Lipid peroxidation levels measured by BODIPY C11 staining in different human cell lines after treatment with Romidepsin (3 nM), MS1943 (5 µM), Tazemetostat (5 µM) or a combination for 24h. I) ROS levels in T-ALL PDX model XAB22 after *ex vivo* treatment with Romidepsin (1 nM), MS1943 (5 µM), Tazemetostat (5 µM) or a combination for 24h. J) Lipid peroxidation levels measured by BODIPY C11 staining in T-ALL PDX model XAB22 after *ex vivo* treatment with Romidepsin (1 nM), MS1943 (5 µM), Tazemetostat (5 µM) or a combination for 24h. K) Lipid peroxidation levels measured by BODIPY C11 staining in AML PDX model XAA2 after *ex vivo* treatment with Romidepsin (2 nM), MS1943 (5 µM), Tazemetostat (5 µM) or a combination for 24h. L) RPMI-8402 cells treated with Romidepsin (3 nM), MS1943 (5 µM), Tazemetostat (5 µM) or a combination for 48h in the presence or absence of 5 mM Gluthation or 10 µM Ferrostatin-1.

## Discussion

Epigenetic deregulation is a hallmark of many hematological malignancies and is a driving force for cellular transformation. In this study we focused on the epigenetic factor EZH2, which was once primarily recognized as the methyltransferase component of the PRC2 complex. However, recent findings describe that EZH2 has a partially disordered transactivation domain^21^ which can be unlocked via phosphorylation of EZH2 on specific residues. EZH2 phosphorylation also induces a release from the PRC2 complex^1,2,22^, which allows it to bind directly to promoter regions and act as a transcriptional regulator in cooperation with MYC transcription factors in different hematological malignancies^1,3,4^.

Since classical EZH2 enzymatic inhibitors only target the methyltransferase activity and not the transcriptional activation of EZH2, we first sought to investigate the efficacy of recently developed EZH2 degraders. Indeed, most of the human hematological cancer cell lines that we tested were only mildly sensitive to EZH2 enzymatic inhibition, even over a longer period of time. Conversely, EZH2 degradation more strongly decreased cellular proliferation and induced higher levels of apoptosis in both cell lines and PDX models of different types of hematological cancers. Notably, we did not have a B cell lymphoma cell line available that harbors an EZH2 methyltransferase activating mutation, such as the EZH2^Y641F^ alteration. We expect that these cells would be sensitive to both EZH2 degradation and EZH2 enzymatic inhibition, as they are specifically addicted to this mutation to sustain their proliferation^23,24^.

To investigate why EZH2 degradation is more powerful than EZH2 enzymatic inhibition in inducing cell death, we performed a transcriptomic analysis after treatment with both types of drugs. While EZH2 degradation led to a swift downregulation of non-canonical EZH2 target genes, canonical EZH2 target genes were only mildly affected. Most likely, EZH2 degradation prevents the direct association of EZH2 with its non-canonical target genes, thereby reducing transcriptional activation. In contrast, inhibition of the PRC2 complex via targeting the enzymatic activity of EZH2 does not immediately lead to changes in gene expression, as its effect on gene activity occurs indirectly via the modification of histone marks. Therefore, we prolonged the treatment with the EZH2 enzymatic inhibitor until we could observe a marked reduction in H3K27me3 levels before performing the transcriptomic analysis. We observed that enzymatic inhibition of EZH2 induced upregulation of canonical EZH2 target genes, as expected, but it also induced upregulation of non-canonical EZH2 target genes. This could potentially be explained by the binding of the bulky inhibitor to EZH2, which reduces its binding affinity with other PRC2 complex members. This may result in an excess of non-PRC2-bound EZH2 that subsequently acts as transcriptional activator of its non-canonical target genes. We also observed that EZH2 degradation induced stronger upregulation of genes implicated in fatty acid and cholesterol synthesis compared to EZH2 enzymatic inhibition. In line with this, the role of EZH2 in the regulation of cholesterol synthesis was recently also described in head and neck squamous cell carcinoma^25^, mesothelioma^26^ and murine melanoma^27^.

We previously identified a striking synergy between EZH2 degradation and HDAC inhibition in a murine model of MYCN-driven T cell lymphoma^1^. We now confirm that this treatment combination is effective in a broader spectrum of hematological malignancies, and that HDAC inhibitors can prompt cells sensitivity to EZH2 enzymatic inhibition. The upregulation of the fatty acids and cholesterol biosynthesis pathways as a result of EZH2 targeting is of particular importance for this synergistic effect. First of all, the cholesterol biosynthesis pathway heavily relies on the availability of acetyl-CoA, as this is the unique carbon source used in this pathway. Histone acetylation, induced by the inhibition of HDACs, also solely relies on acetyl-CoA as acetyl-donor. Furthermore, SAT1 hyperactivity has also been shown to deplete acetyl-CoA levels, as this enzyme catalyzes the formation of N-acetylspermidine/N-acetylspermine from the polyamines spermidine/spermine^28^. Consequently, the acetyl-CoA levels needed to fuel the TCA cycle to generate cellular energy or important intermediary metabolites are lacking, and the cells experience metabolic stress due to the combination treatment. Furthermore, increased fatty acid and cholesterol biosynthesis leads to depletion of antioxidants in the cell, thereby increasing the cellular ROS levels. Combined with a higher cellular lipid content, this leads to the induction of lipid peroxidation, a process in which oxidized phospholipids accumulate, leading to the disruption of plasma membrane integrity and resulting in ferroptotic cell death^29^. Previous studies have already shown that HDAC inhibition shows synergistic effects with EZH2 enzymatic inhibition in germinal-center-derived lymphoma^11^, multiple myeloma^12^ or non-small cell lung cancer^10^, but our study is the first to demonstrate this effect in several types of hematological malignancies, while also uncovering the molecular mechanism behind the synergy and showing that ferroptosis is the main mechanism of cell death.

Our findings on the molecular mechanism underlying EZH2 targeting and its synergy with HDAC inhibition have strong implications for future clinical applications. EZH2 degradation suggests to be a more powerful approach for effective growth inhibition compared to EZH2 enzymatic inhibition, at least for hematological cancers that are not characterized by hyperactivation of the methyltransferase activity of EZH2. As many hematological malignancies are characterized by high levels of EZH2, this treatment approach could be broadly applied in both lymphoid and myeloid malignancies. Furthermore, FDA-approved HDAC inhibitors are associated with significant toxicities and a limited efficacy. Combined targeting of EZH2 and HDACs could improve their effect on growth inhibition, while reducing the associated toxicities. As the synergistic effect of combined EZH2 and HDAC targeting is also observed with a clinically approved EZH2 enzymatic inhibitor, translation to the clinic could be swift. Finally, our insights in the molecular mechanism underlying the synergistic effect of combined EZH2 and HDAC targeting could lead to the development of early biomarkers of response. In conclusion, combined EZH2 and HDAC targeting is a promising therapeutic approach for many different hematological malignancies.

## Methods

### Cell lines

All cell lines (**Table S1**) were obtained from DSMZ. The cells were cultured in RPMI 1640 medium supplemented with 20% fetal calf serum (Invitrogen).

### Patient-derived xenograft (PDX) models

Informed consent was obtained from all subjects. PDX characteristics are shown in **Table S2**. All PDX models were propagated in in-house bred NSG or NSG-SGM3 mice. Mice were housed in individually ventilated cages with a temperature between 18-23°C and humidity between 40-60% in SPF or semi-SPF conditions in the KU Leuven animal facility. In the case of T-ALL or B-ALL PDX samples, 1×10^6^ cells were injected per NSG mouse. For AML PDX samples, 5×10^6^ cells were injected per NSG-SGM3 mouse. Disease development was followed by biweekly blood withdrawal. The peripheral blood was submitted to red blood cell lysis, followed by hCD45 staining (Biolegend) and detected using flow cytometry (MACSQuant VYB (Miltenyi)). NSG or NSG-SGM3 mice that presented over 50% of hCD45 in the peripheral blood were euthanized.

For the *ex vivo* experiments, the spleen was collected from moribund PDX mice, and processed to obtain a single-cell suspension. Red blood cells were removed using a red blood cell lysis buffer (150 nM NH_4_Cl, 0.1mM EDTA, 10mM KHCO_3_ dissolved in 1 L of MiliQ water).

Mouse experiments were approved and supervised by the KU Leuven ethical committee and conducted according to EU legislation (Directive 2010/63/EU).

### *Ex vivo / In vitro* small molecule treatments

All small molecule compounds used in this study are listed in **Table S3**.

The cells (cell lines or a single-cell suspension from a PDX model) were seeded into 12-well, 24-well or 96-well plates at a concentration of 5×10^5^ cells/mL in RPMI 1640 supplemented with 20% fetal calf serum and 1X primocin (Invivogen).

For 12-well and 24-well plates, the compounds (all dissolved in DMSO) were added and DMSO concentration was normalized. Cell proliferation was measured after 24h, 48h or 72h using a MACSQuant VYB (Miltenyi).

For 96-well plates, the compounds (all dissolved in DMSO) were added in a randomized fashion using a D300e digital dispenser (Tecan) and the DMSO concentration was normalized. Cell proliferation was measured after 48h or 72h using the ATPlite luminescence system (PerkinElmer) with a Victor X4 multilabel plate reader.

### Flow Cytometry

Annexin V / PI stainings were performed using the APC/FITC Annexin V Apoptosis Detection Kit (BioLegend) according to the manufacturer’s instructions. Intracellular stainings were performed using the Foxp3 / Transcription Factor Staining kit (Thermo Fischer Scientific) according to the manufacturer’s instructions. Antibodies are listed in **Table S4**.

Cell cycle analysis was performed via DAPI staining (100 ng/mL).

ROS levels were measured using H2DCFDA (1h incubation at a concentration of 5 µM) or using a ROS assay Kit (Thermo Fischer Scientific) according to the manufacturer’s instructions. Lipid peroxidation levels were measured using BODIPY C11 (1h incubation at a concentration of 2 µM).

All flow cytometric data was acquired on a MACSQuant VYB (Miltenyi) or on a Fortessa X-20 (BD Biosciences). Data were analyzed with the FlowJo software (Tree Star).

### Western Blotting

Cell lysates were prepared using 1X Cell Lysis Buffer (Cell Signaling Technologies) containing protease inhibitor (Complete – EDTA-free, Roche) and 1 mM Na_3_VO_4_. Proteins were separated by SDS-PAGE (NuPAGE NOVEX Bis-Tris 4–12% gels (Life Technologies) and transferred to PVDF membranes. Subsequent labelling was carried out using unlabeled primary antibodies. Western blot detection was performed with secondary antibodies conjugated with horseradish peroxidase (rabbit (GE Healthcare) or mouse (Sigma-aldrich)). Images were acquired using a cooled charge-coupled device camera system (Vilber, Fusion FX). Antibodies used for Western Blotting are listed in **Table S5**.

### RNA extraction

RNA from mouse tissue and from cell lines was extracted using the Maxwell RSC simplyRNA Cells Kit (Promega) on the Maxwell RSC instrument according to the manufacturer’s instructions.

### cDNA synthesis and qRT-PCR

cDNA synthesis was carried out using GoScript (Promega) and qRT-PCR was performed using the GoTaq qRT-PCR master mix (Promega) with the ViiA7 Real Time PCR system (Applied Biosystems). Primers used for qRT-PCR are listed in **Table S6**.

For qRT-PCR analyses, data are expressed as the mean ± standard deviation (SD). Comparisons between two groups were performed using one-way ANOVA with corrections for multiple comparisons.

### ChIPmentation ChIP-sequencing

ChIPmentation ChIP-seq was performed as described previously^30–32^. 20-50 million primary spleen cells were washed with PBS and cross-linked with 1% formaldehyde for 10 min and then quenched by addition of glycine. For nuclei isolation, cells were resuspended in 1X RSB buffer (10 mM Tris pH7.4, 10 mM NaCl, 3 mM MgCl2) and left on ice for 10 min. Cells were collected and resuspended in RSBG40 buffer (10 mM Tris pH7.4, 10 mM NaCl, 3 mM MgCl2, 10% glycerol, 0.5% NP40) with 1/10 v/v of 10% detergent (3.3% w/v sodium deoxycholate, 6.6% v/v Tween-40). Nuclei were resuspended in L3B buffer (10 mM Tris-Cl pH 8.0, 100 mM NaCl, 1 mM EDTA, 0.5 mM EGTA, 0.1% Na-Deoxycholate, 0.5% N-Lauroylsarcosine, 0.2% SDS). Chromatin was fragmented to 200-400 bp using a Bioruptor (Diagenode) for 20-25 cycles (30 s on, 30 s off, High). The chromatin was supplemented with 1% Triton-×100 after fragmentation. The antibody (H3K27ac, Abcam ab4729) was pre-conjugated to magnetic protein A/G beads (Millipore). Chromatin immunoprecipitation was carried out overnight. Tagmentation and library preparation was performed using the Nextera DNA library prep kit (Illumina). DNA was purified using triple sided SPRI bead clean-up (Agencourt AMPure Beads, Beckman Coulter).

### Next generation sequencing and data analysis

RNA and ChIPmentation libraries were sequenced on a HiSeq 4000 with 125bp single-end reads (Illumina). All data was first cleaned with fastq-mcf from ea-utils, after which a quality control was performed with FastQC. Subsequently the reads were mapped to the respective reference genomes (GRCh38/hg38) with HISAT2. Further processing of the reads was executed with the SAMtools package and the number of reads per transcript was determined with HTSeq-count. DESeq2 was used to determine the significantly differentially expressed genes. Heatmaps and expression plots were constructed with R. Gene set enrichment analyses were performed with the GSEA software from the Broad Institute, with the genes ranked according to the following score; -sign(log2FoldChange)*log(p_adj). Possible regulatory transcription factor motifs were determined with i-CisTarget^33^.

ChIPmentation sequencing data was preprocessed similarly as the RNA-seq data, i.e. cleaning with fastq-mcf and quality control with FastQC. Subsequently, these cleaned reads were mapped to the respective reference genome (GRCh38/hg38) with the HISAT2 software, after which the reads were further processed with SAMtools. The ChIP signals were normalized with the deepTools package (bamCoverage) and the peaks were called with MACS2. These peaks were then annotated with a custom script. The centered heatmaps were constructed with the deepTools package (computeMatrix, plotHeatmap).

### Statistical analysis

Unless stated otherwise, experiments were performed at least three times independently and graphs represent mean +/- standard deviation. All statistical analyses were performed using Prism software (Graphpad software, CA, USA), using unpaired t-tests or on-way or two-way ANOVA with Tukey’s multiple comparison correction.

## Data availability

The RNA-seq and ChIP-seq data were deposited in the Gene Expression Omnibus database with accession number GSE254725.

## Acknowledgements

The authors thank the KU Leuven Genomics Core and the VIB FACS expertise center for their technical services. AV is supported by a postdoctoral fellowship from the Research Foundation Flanders (FWO), MVB and SD are supported by postdoctoral mandates for fundamental research from the Foundation against Cancer (Stichting Tegen Kanker). This study was further supported by the Fund Tom Debackere for Lymphoma research.

## Author Contributions

AV and MVB designed the study, collected and analyzed data, and wrote the manuscript. SD performed bioinformatic analyses. TS, YS, JS, SVB, MA, KA, N Meeuws, IB, FR, LB, N Mentens, OG and KJ collected and analyzed data. MA, KA, LB, OG and HS collected patient samples for PDX model generation. TS, SD and YS edited the manuscript. JC, DD and MVB supervised the study.

## References

1. Vanden Bempt, M., et al. Aberrant MYCN expression drives oncogenic hijacking of EZH2 as a transcriptional activator in peripheral T cell lymphoma. Blood 140, 2463–2476 (2022).

2. Yan, J. et al. EZH2 phosphorylation by JAK3 mediates a switch to noncanonical function in natural killer/T-cell lymphoma. Blood 128, 948–58 (2016).

3. Wang, J. et al. EZH2 noncanonically binds cMyc and p300 through a cryptic transactivation domain to mediate gene activation and promote oncogenesis. Nat. Cell Biol. 3, 384–399 (2022).

4. Yu, X. et al. Dissecting and targeting noncanonical functions of EZH2 in multiple myeloma via an EZH2 degrader. Oncogene 42, 994–1009 (2023).

5. Ma, A. et al. Discovery of a first-in-class EZH2 selective degrader. Nat. Chem. Biol. 16, 214–222 (2020).

6. Wang, C. et al. Discovery of precision targeting EZH2 degraders for triple-negative breast cancer. Eur. J. Med. Chem. 238, 114462 (2022).

7. Stubbs, M. C. et al. Selective Inhibition of HDAC1 and HDAC2 as a Potential Therapeutic Option for B-ALL. Clin. Cancer Res. 21, 2348–2358 (2015).

8. Lu, G. et al. Update on histone deacetylase inhibitors in peripheral T-cell lymphoma (PTCL). Clin. Epigenetics 15, 124 (2023).

9. Schade, A. E. et al. Combating castration-resistant prostate cancer by co-targeting the epigenetic regulators EZH2 and HDAC. PLoS Biol. 21, e3002038 (2023).

10. Takashina, T. et al. Combined inhibition of EZH 2 and histone deacetylases as a potential epigenetic therapy for non-small-cell lung cancer cells. Cancer Sci. 107, 955–962 (2016).

11. Lue, J. K. et al. Precision Targeting with EZH2 and HDAC Inhibitors in Epigenetically Dysregulated Lymphomas. Clin. Cancer Res. 25, 5271–5283 (2019).

12. Harding, T. et al. EZH2 inhibitors sensitize myeloma cell lines to panobinostat resulting in unique combinatorial transcriptomic changes. Oncotarget 9, 21930–21942 (2018).

13. Kamminga, L. M. et al. The Polycomb group gene Ezh2 prevents hematopoietic stem cell exhaustion. Blood 107, 2170–9 (2006).

14. Huang, Y. et al. Characterizing cancer metabolism from bulk and single-cell RNA-seq data using METAFlux. Nat. Commun. 14, 4883 (2023).

15. Guertin, D. A. et al. Acetyl-CoA metabolism in cancer. Nat. Rev. Cancer 23, 156–172 (2023).

16. Ou, Y. et al. Activation of SAT1 engages polyamine metabolism with p53-mediated ferroptotic responses. Proc. Natl. Acad. Sci. U. S. A. 113, E6806–E6812 (2016).

17. Doll, S. et al. ACSL4 dictates ferroptosis sensitivity by shaping cellular lipid composition. Nat. Chem. Biol. 13, 91–98 (2017).

18. Freitas, F. P. et al. 7-Dehydrocholesterol is an endogenous suppressor of ferroptosis. Nature 626, 401– 410 (2024).

19. Li, Y. et al. 7-Dehydrocholesterol dictates ferroptosis sensitivity. Nature 626, 411–418 (2024).

20. Stockwell, B. R. et al. Ferroptosis: A Regulated Cell Death Nexus Linking Metabolism, Redox Biology, and Disease. Cell 171, 273–285 (2017).

21. Jiao, L. et al. A partially disordered region connects gene repression and activation functions of EZH2. Proc. Natl. Acad. Sci. U. S. A. 117, 16992–17002 (2020).

22. Xu, K. et al. EZH2 oncogenic activity in castration-resistant prostate cancer cells is Polycomb-independent. Science 338, 1465–9 (2012).

23. Souroullas, G. P. et al. An oncogenic Ezh2 mutation induces tumors through global redistribution of histone 3 lysine 27 trimethylation. Nat. Med. 22, 632–40 (2016).

24. Yap, D. B. et al. Somatic mutations at EZH2 Y641 act dominantly through a mechanism of selectively altered PRC2 catalytic activity, to increase H3K27 trimethylation. Blood 117, 2451–2459 (2011).

25. Xu, X. et al. Targeting epigenetic modulation of cholesterol synthesis as a therapeutic strategy for head and neck squamous cell carcinoma. Cell Death Dis. 12, 482 (2021).

26. Pandey, G. K. et al. Genetic screens reveal new targetable vulnerabilities in BAP1-deficient mesothelioma. Cell reports. Med. 4, 100915 (2023).

27. Zhang, T. et al. Dysregulated lipid metabolism blunts the sensitivity of cancer cells to EZH2 inhibitor. EBioMedicine 77, 103872 (2022).

28. Pegg, A. E. Spermidine/spermine- *N*^1^ -acetyltransferase: a key metabolic regulator. Am. J. Physiol. Metab. 294, E995–E1010 (2008).

29. Liang, D. et al. Ferroptosis at the intersection of lipid metabolism and cellular signaling. Mol. Cell 82, 2215–2227 (2022).

30. Vanden Bempt, M., et al. Cooperative Enhancer Activation by TLX1 and STAT5 Drives Development of NUP214-ABL1/TLX1-Positive T Cell Acute Lymphoblastic Leukemia. Cancer Cell 34, 271–285.e7 (2018).

31. de Bock, C. E. et al. HOXA9 Cooperates with Activated JAK/STAT Signaling to Drive Leukemia Development. Cancer Discov. 8, 616–631 (2018).

32. Broux, M. et al. Suz12 inactivation cooperates with JAK3 mutant signaling in the development of T-cell acute lymphoblastic leukemia. Blood 134, 1323–1336 (2019).

33. Verfaillie, A. et al. iRegulon and i-cisTarget: Reconstructing Regulatory Networks Using Motif and Track Enrichment. Curr. Protoc. Bioinforma. 52, 2.16.1–2.16.39 (2015).

