## Supplemental information for "Dual targeting of EZH2 and Histone Deacetylases in hematological malignancies promotes transcriptional and metabolic deregulation leading to ferroptosis"

### Supplemental figures and legends

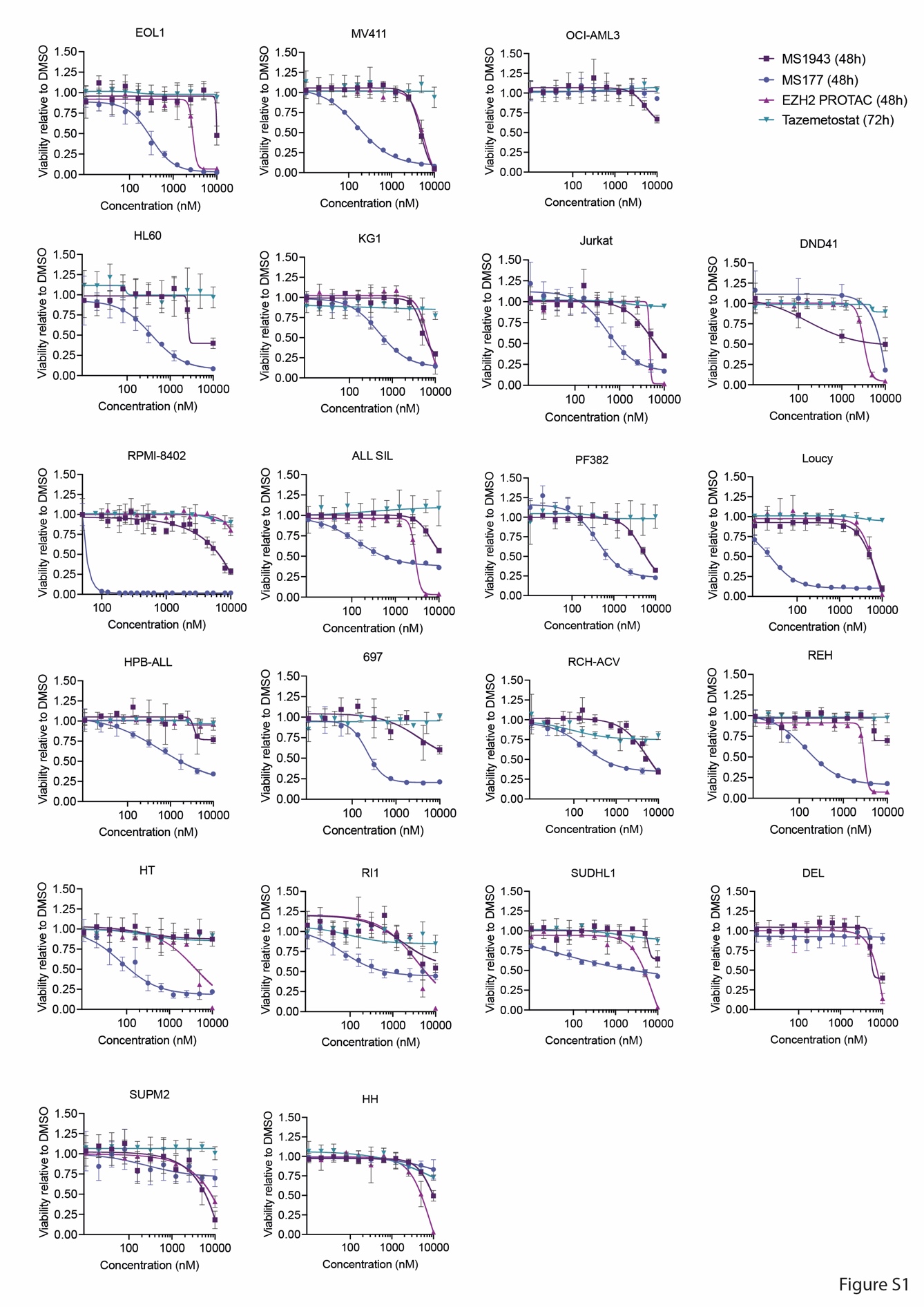

#### Figure S1: Dose response curves for EZH2 degraders and the EZH2 enzymatic inhibitor Tazemetostat in human cell lines of lymphoid and myeloid origin.

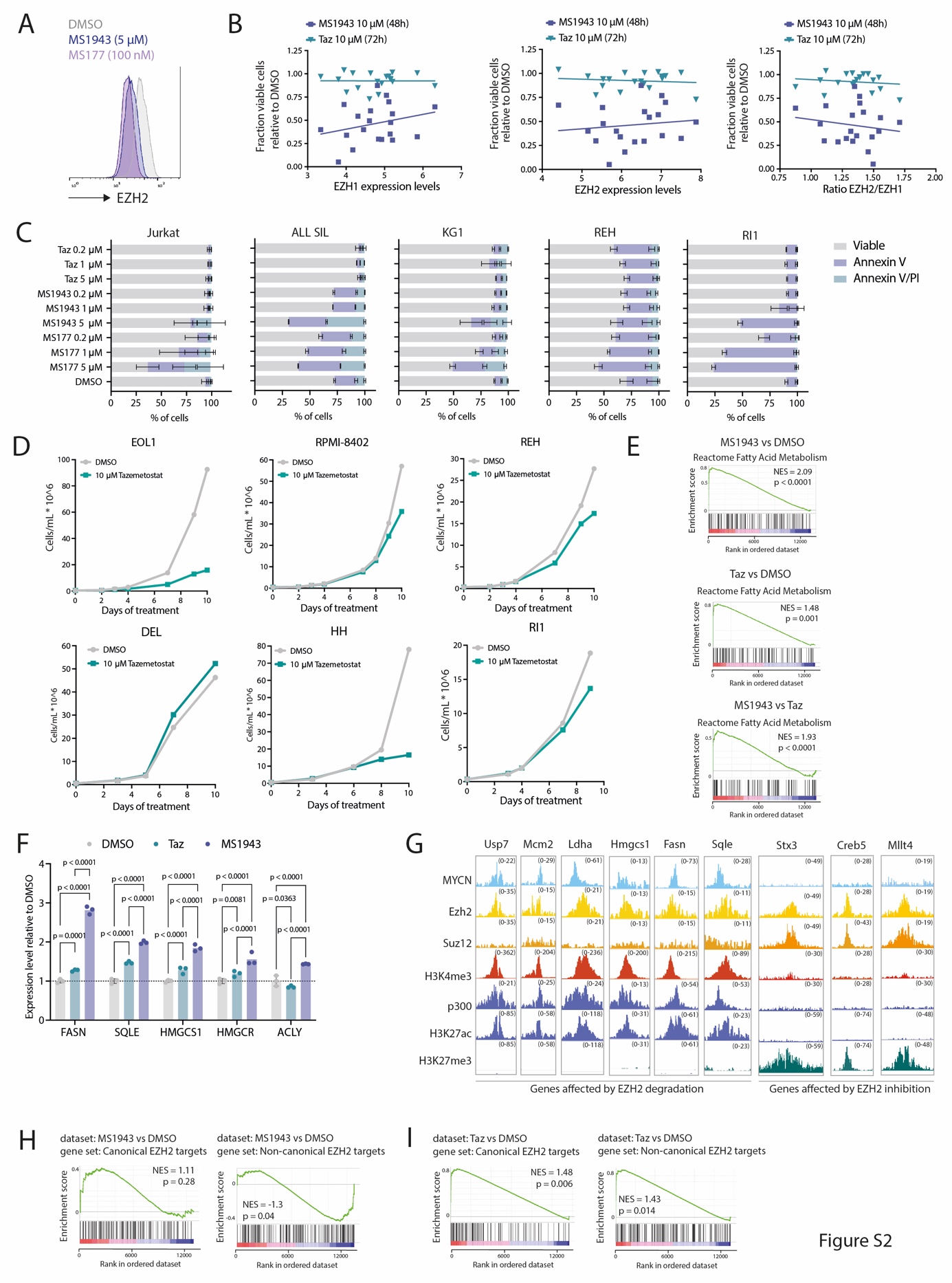

#### Figure S2: Both myeloid and lymphoid malignancies are sensitive to EZH2 degradation.

A) Intracellular staining for EZH2 in RPMI-8402 cells treated with EZH2 degraders (MS1943 or MS177) for 16h at the indicated concentrations.
B) Correlation between EZH1/2 expression (left and middle) or the ratio of EZH2/EZH1 expression (right) with the response to EZH2 degradation or EZH2 enzymatic inhibition at the indicated concentrations in human cell lines. Expression levels are exported from the publicly available DepMap portal.
C) Annexin V / PI staining of the Jurkat, ALL SIL, KG1, REH and RI1 cell lines after 48h treatment with an EZH2 enzymatic inhibitor (Tazemetostat) or EZH2 degraders (MS177, MS1943) at different concentrations.
D) Growth curve of the EOL1, RPMI-8402, REH, DEL, HH and RI1 cell lines treated with 10 µM Tazemetostat or DMSO for 10 days.
E) Gene set enrichment analysis (GSEA) on the differentially expressed genes in RPMI-8402 cells treated with 5 µM MS1943 for 16h compared to DMSO (top), 5 µM Tazemetostat for 72h compared to DMSO (middle) or 5 µM MS1943 for 16h compared to 5 µM Tazemetostat for 72h (bottom).
F) qRT-PCR analysis of cholesterol biosynthesis gene expression in EOL1 cells treated with 5 µM MS1943 or 5 µM Tazemetostat for 24h.
G) Representative ChIP-seq tracks for the indicated promoter regions showing binding of MYCN, Ezh2, Suz12, p300, and histone marks H3K4me3, H3K27ac, and H3K27me3 in murine MYCN-driven TCL^1^.
H) Gene set enrichment analysis (GSEA) on the differentially expressed genes in RPMI-8402 cells treated with 5 µM MS1943 for 16h compared to DMSO. The gene sets used for this analysis are the top-250 genes with EZH2 + H3K27me3 peaks (canonical EZH2 targets) or the top-250 genes with EZH2 + H3K27ac peaks (non-canonical EZH2 genes) as determined in murine MYCN-driven T cell lymphoma^1^.
I) Gene set enrichment analysis (GSEA) on the differentially expressed genes in RPMI-8402 cells treated with 5 µM Tazemetostat for 72h compared to DMSO. The gene sets used for this analysis are the top-250 genes with EZH2 + H3K27me3 peaks (canonical EZH2 targets) or the top-250 genes with EZH2 + H3K27ac peaks (non-canonical EZH2 genes) as determined in murine MYCN-driven T cell lymphoma^1^.

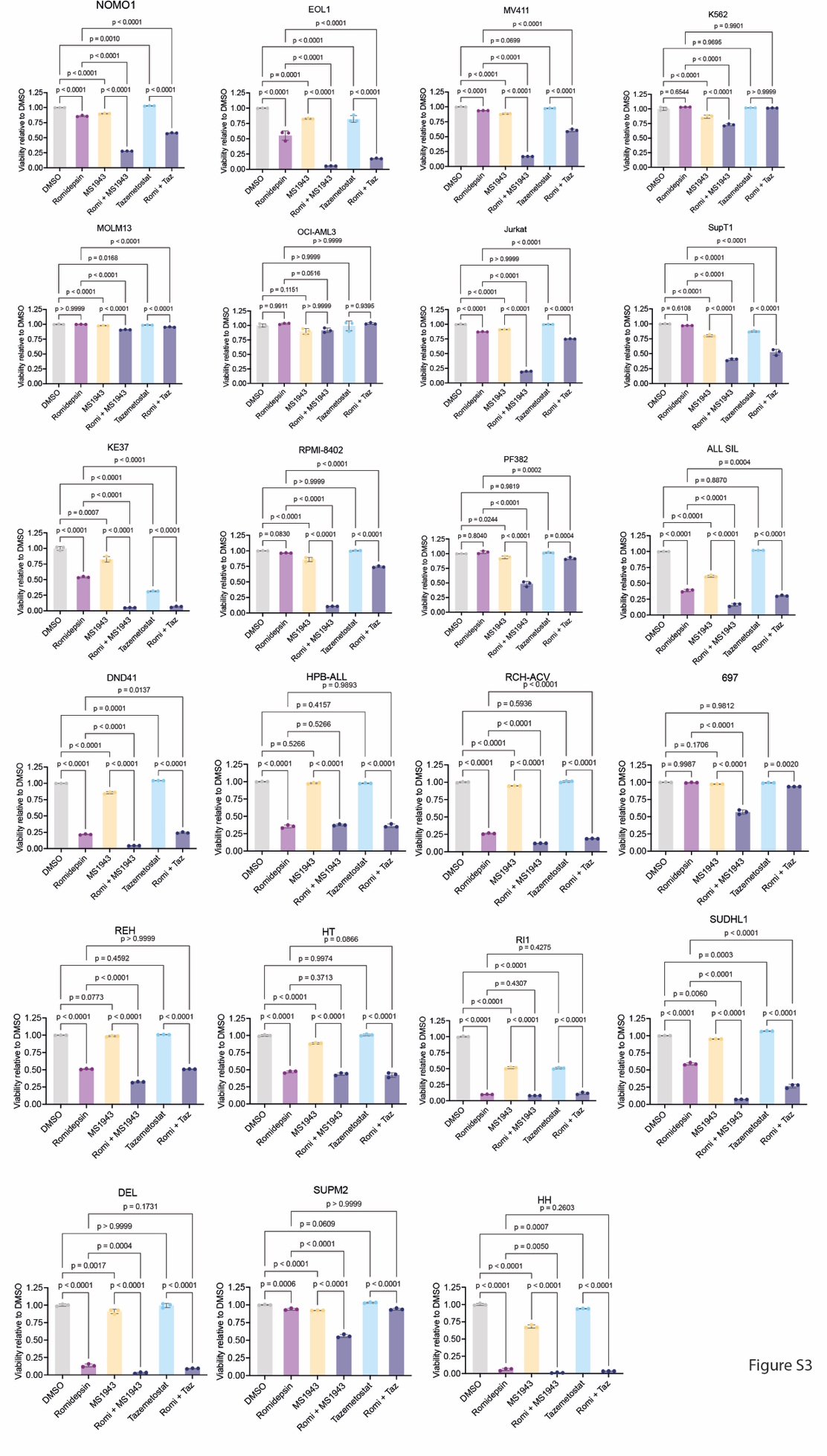

#### Figure S3: Synergistic effects between EZH2 inhibition/degradation and HDAC inhibition.

All cell lines are treated for 48h with Romidepsin (3 nM), MS1943 (5 µM), Tazemetostat (5 µM) or a combination. Viability was measured on a MacsQuant VYB Flow Cytometer.

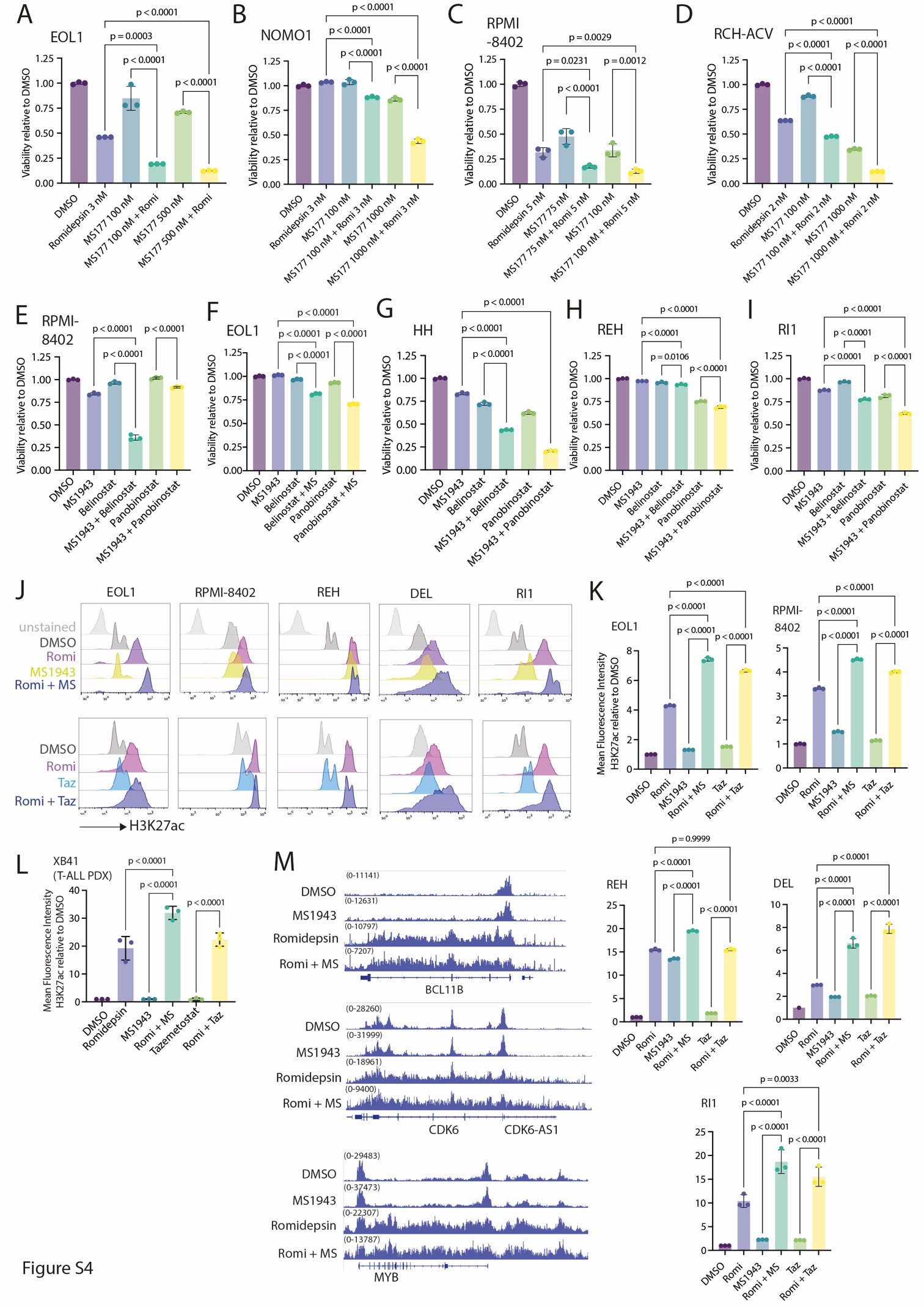

#### Figure S4: Dual targeting of EZH2 and HDACs induces cell death in myeloid and lymphoid malignancies.

A) Viability of EOL1 cells treated for 18h with Romidepsin and/or MS177 at the indicated concentrations.
B) Viability of NOMO1 cells treated for 48h with Romidepsin and/or MS177 at the indicated concentrations.
C) Viability of RPMI-8402 cells treated for 48h with Romidepsin and/or MS177 at the indicated concentrations.
D) Viability of RCH-ACV cells treated for 72h with Romidepsin and/or MS177 at the indicated concentrations.
E) Viability of RPMI-8402 cells treated for 48h with Belinostat (90 nM), Panobinostat (1 nM), MS1943 (5 µM) or a combination.
F) Viability of EOL1 cells treated for 48h with Belinostat (90 nM), Panobinostat (1 nM), MS1943 (5 µM) or a combination.
G) Viability of HH cells treated for 48h with Belinostat (250 nM), Panobinostat (15 nM), MS1943 (5 µM) or a combination.
H) Viability of REH cells treated for 48h with Belinostat (100 nM), Panobinostat (8 nM), MS1943 (5 µM) or a combination.
I) Viability of RI1 cells treated for 48h with Belinostat (100 nM), Panobinostat (8 nM), MS1943 (5 µM) or a combination.
J) Intracellular H3K27ac staining of EOL1, RPMI-8402, REH, DEL and RI1 cell lines treated with Romidepsin (3 nM), MS1943 (5 µM), Tazemetostat (5 µM) or a combination.
K) Quantification of the mean fluorescent intensity of the H3K27ac signal in the EOL1, RPMI-8402, REH, DEL and RI1 cell lines treated with Romidepsin (3 nM), MS1943 (5 µM), Tazemetostat (5 µM) or a combination.
L) Quantification of the mean fluorescent intensity of the H3K27ac signal in *ex vivo* treated cells from the XB41 T-ALL PDX model treated with Romidepsin (1 nM), MS1943 (5 µM), Tazemetostat (5 µM) or a combination for 24h.
M) H3K27ac ChIP-seq tracks for *BCL11b*, *CDK6* and *MYB* in RPMI-8402 cells treated with DMSO, Romidepsin (3 nM), MS1943 (5 µM) or Romidepsin + MS1943 for 16h.

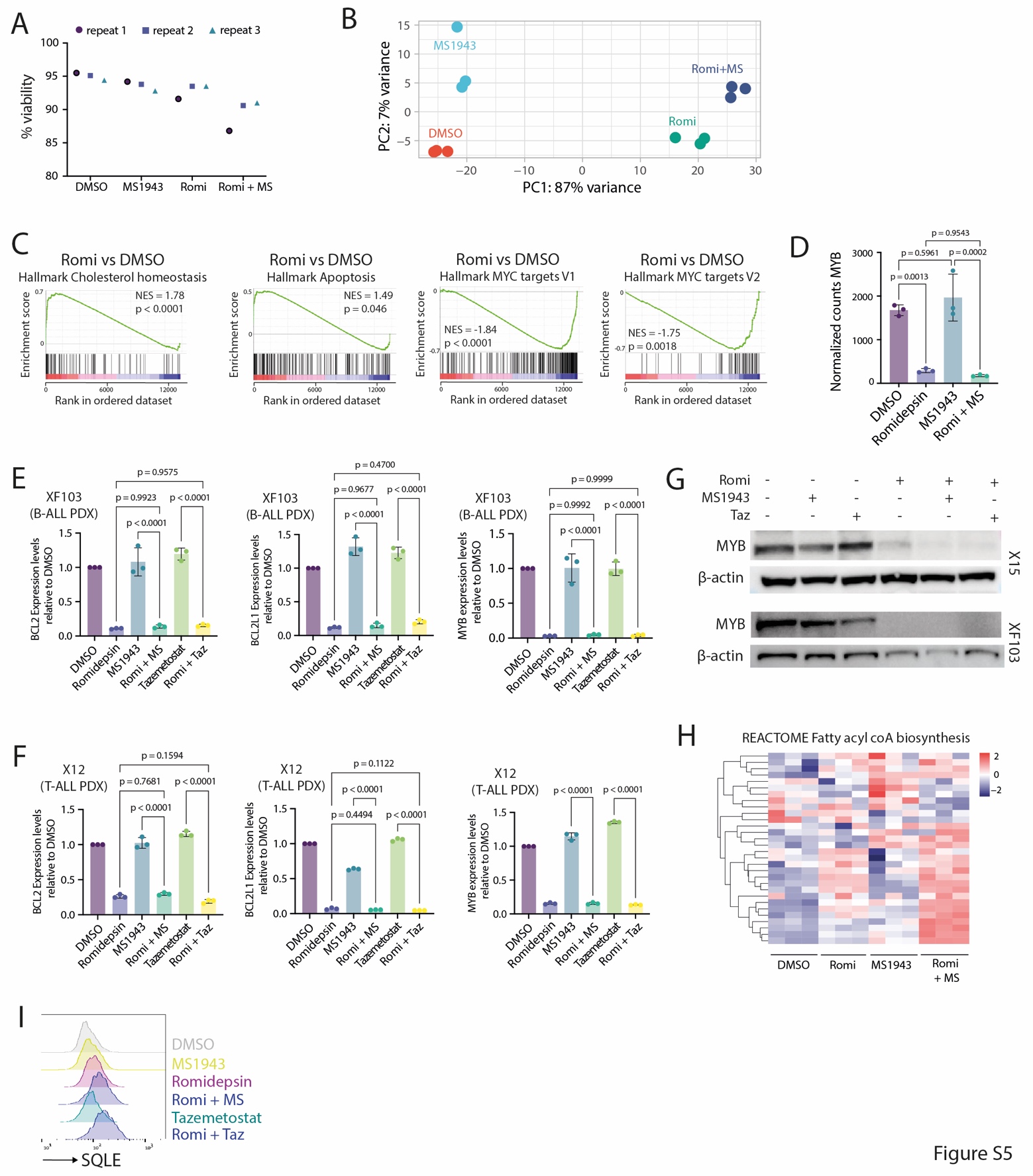

#### Figure S5: Combined targeting of EZH2 and HDAC induces activation of cholesterol biosynthesis.

A) Viability at the time of RNA extraction of RPMI cells treated with DMSO, Romidepsin (3 nM), MS1943 (5 µM) or Romidepsin + MS1943 for 16h (samples used for RNA-sequencing).
B) Principal component analysis of RNA-sequencing data from RPMI-8402 cells treated with DMSO, Romidepsin (3 nM), MS1943 (5 µM) or Romidepsin + MS1943 for 16h.
C) Gene set enrichment analysis (GSEA) on the differentially expressed genes in RPMI-8402 cells treated with Romidepsin (3 nM) compared to DMSO for 16h.
D) Normalized counts of *MYB* in RPMI-8402 cells treated with DMSO, Romidepsin (3 nM), MS1943 (5 µM) or the combination for 16h.
E) qRT-PCR analysis of gene expression levels in XF103 B-ALL PDX cells treated *ex vivo* with DMSO, Romidepsin (3 nM), Tazemetostat (5 µM), MS1943 (5 µM) or the combination for 24h.
F) qRT-PCR analysis of gene expression levels in X12 T-ALL PDX cells treated *ex vivo* with DMSO, Romidepsin (3 nM), Tazemetostat (5 µM), MS1943 (5 µM) or the combination for 24h.
G) Western Blot showing expression of MYB in X15 T-ALL or XF103 B-ALL PDX cells treated *ex vivo* with DMSO, Romidepsin (5 nM), MS1943 (5 µM) or the combination for 24h.
H) Heatmap showing the expression levels of the genes in the REACTOME Fatty acyl coA biosynthesis gene set in RPMI-8402 cells treated with DMSO, Romidepsin (3 nM), MS1943 (5µM), or the combination for 16h.
I) Intracellular staining showing SQLE levels in RPMI-8402 cells treated with DMSO, Romidepsin (3 nM), MS1943 (5 µM) or the combination for 16h.

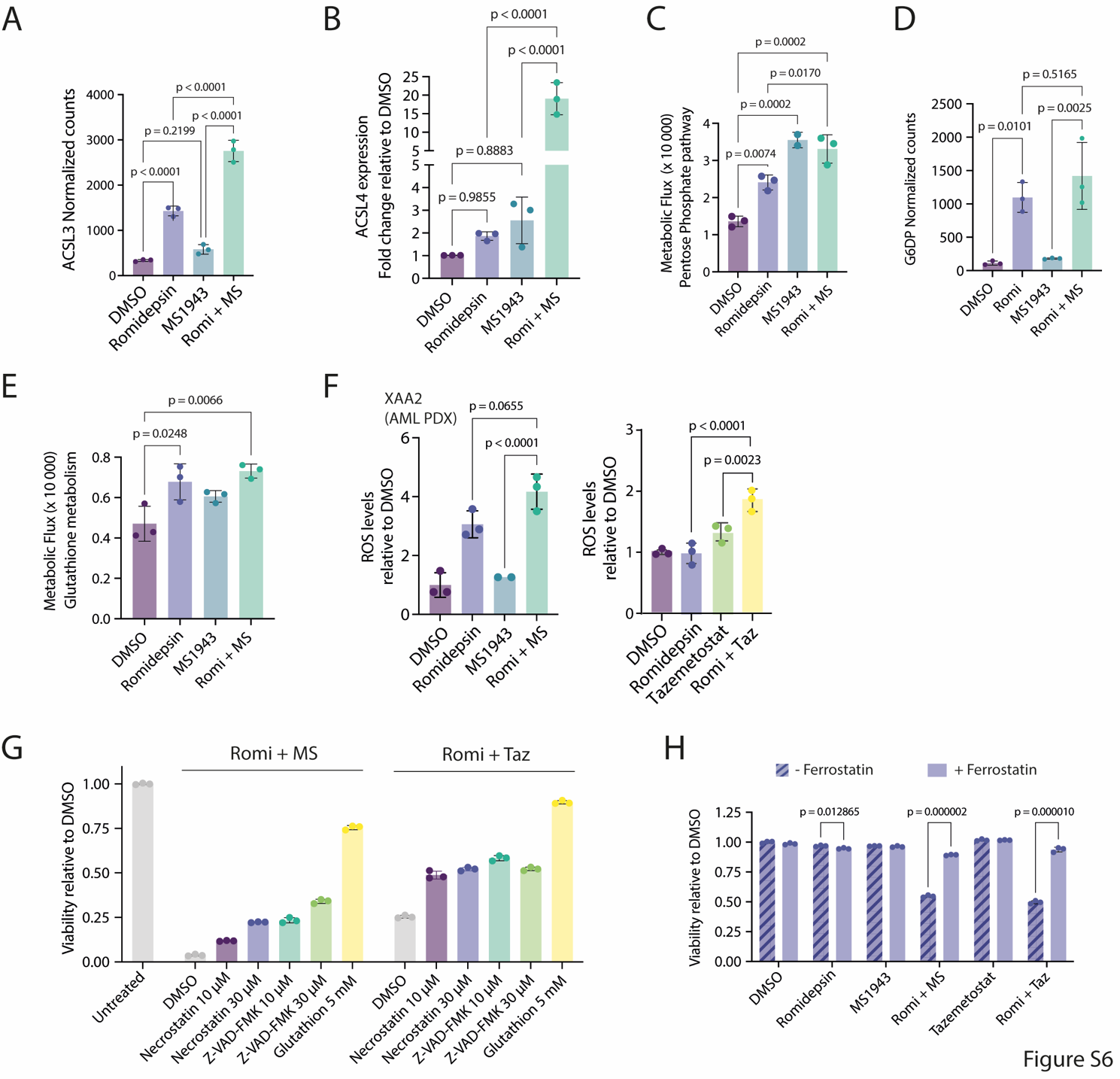

#### Figure S6: Combined targeting of EZH2 and HDAC leads to ferroptosis induction.

A) Normalized counts of *ACSL3* in RPMI-8402 cells treated with DMSO, Romidepsin (3 nM), MS1943 (5 µM) or the combination for 16h.
B) qRT-PCR analysis of ACSL4 expression levels in RPMI-8402 cells treated with DMSO, Romidepsin (3 nM), MS1943 (5 µM) or the combination for 16h.
C) Pathway activation scores (~Metabolic flux) of the pentose phosphate pathway as inferred using the METAFlux R package in RPMI-8402 cells treated with DMSO, Romidepsin (3 nM), MS1943 (5 µM) or the combination for 16h.
D) Normalized counts of *G6DP* (Glucose-6-phosphate dehydrogenase, the rate-limiting enzyme in the pentose phosphate pathway) in RPMI-8402 cells treated with DMSO, Romidepsin (3 nM), MS1943 (5 µM) or the combination for 16h.
E) Pathway activation scores (~Metabolic flux) of glutathione metabolism as inferred using the METAFlux R package in RPMI-8402 cells treated with DMSO, Romidepsin (3 nM), MS1943 (5 µM) or the combination for 16h.
F) ROS levels in AML PDX model XAA2 after *ex vivo* treatment with Romidepsin (2 (left) or 1 (right) nM), MS1943 (5 µM), Tazemetostat (5 µM) or a combination for 24h.
G) Viability of RPMI-8402 cells treated for 48h with DMSO, Romidepsin (3 nM), MS1943 (5 µM) or the combination, with or without Necrostatin (necroptosis inhibitor), Z-VAD-FMK (apoptosis inhibitor) or glutathione (ferroptosis inhibitor) at the indicated concentrations.
H) Viability of DEL cells treated for 48h with DMSO, Romidepsin (3 nM), MS1943 (5 µM) or the combination, with or without Ferrostatin-1 (ferroptosis inhibitor).

### Supplemental tables

Table S1: list of cell lines used in this manuscript

| CELL LINE | Disease subtype | Genetic characteristics | Sex |
| --- | --- | --- | --- |
| NOMO1 | AML | MLL-AF9 fusion | F |
| EOL1 | AML | FIP1L1-PDGFRa fusion | M |
| MV411 | AML | MLL-AFF1 fusion  FLT3 ITD | M |
| K562 | CML | BCR-ABL1 fusion | F |
| OCI-AML3 | AML | NRAS Q61L  NPM1 inactivation | M |
| HL60 | AML | NRAS Q61L  TP53 deletion (homozygous) | F |
| MOLM13 | AML | MLL-AF9 fusion  FLT3 ITD | M |
| KG1 | AML | NRAS G12A  FGFR1OP2-FGFR1 fusion | M |
| JURKAT | T-ALL | NOTCH1, FBXW7  TAL1 overexpression | M |
| SUPT1 | T-ALL | PIK3CA | M |
| KE37 | T-ALL | NOTCH1  NRAS G12A  TAL1 overexpression | M |
| DND41 | T-ALL | BCL11B-TLX3 fusion  NOTCH1 | M |
| RPMI-8402 | T-ALL | SIL-TAL1 fusion  LMO1-TRD fusion  NOTCH1, FBXW7 | F |
| ALL SIL | T-ALL | NUP214-ABL1 fusion  NOTCH1  TLX1 overexpression | M |
| PF382 | T-ALL | NOTCH1  TAL1 overexpression | F |
| LOUCY | T-ALL (ETP-ALL) | SET-NUP214 fusion | F |
| HPB-ALL | T-ALL | CBFB-MYLPF fusion  NOTCH1, FBXW7  TLX3 overexpression | M |
| 697 | B-ALL | TCF3-PBX1 fusion | M |
| RCH-ACV | B-ALL | TCF3-PBX1 fusion | F |
| REH | B-ALL | ETV6-RUNX1 fusion | F |
| SUPB15 | B-ALL | BCR-ABL1 fusion | M |
| HT | GC-DLBCL |  | M |
| RI1 | ABC-DLBCL |  | F |
| SUDHL1 | ALCL | NPM1-ALK fusion | M |
| DEL | ALCL | NPM1-ALK fusion | M |
| SUPM2 | ALCL | NPM1-ALK fusion | F |
| HH | CTCL | FOXK2-TP63 fusion | M |

Table S2: PDX models used in this manuscript

| PDX model | Disease subtype | Genetic characteristics | Adult or Pediatric | Sex |
| --- | --- | --- | --- | --- |
| X12 | T-ALL | TLX3+, NUP214-ABL1 fusion | unknown | unknown |
| X15 | T-ALL | TAL1+ | unknown | unknown |
| XAB22 | T-ALL /T-LBL | NOTCH1 mutation | adult | male |
| XB41 | T-ALL | TAL1+, NOTCH1 mutation | pediatric | female |
| XF100 | Pre-T-ALL | TLX3+, NOTCH1 mutation | pediatric | male |
| XC51 | B-ALL | BCR-ABL fusion | pediatric | male |
| XF103 | B-ALL | ETV6-RUNX1 fusion, PAX5 deletion | pediatric | female |
| XAA2 | AML | NPM1 mutation, FLT3-ITD | adult | female |

Table S3: small molecule compounds

| Compound | Supplier | Cat# | Cas# |
| --- | --- | --- | --- |
| MS1943 | MedKoo | 462549 | 2225938-17-8 |
| MS177 | DC Chemicals | DC70618 | 2225938-86-1 |
| EZH2 PROTAC (Compound 150d) | MedChem Express | HY-147525 | 2641601-67-2 |
| Tazemetostat | Selleck Chemicals | S7128 | 1403254-99-8 |
| Romidepsin | AdooQ BioScience | A11920 | 128517-07-7 |
| Panobinostat | MedChem Express | HY-10224 | 404950-80-7 |
| Belinostat | MedChem Express | HY-10225 | 866323-14-0 |
| Hymeglusin | Santa Cruz | sc-203077 | 29066-42-0 |
| Lovastatin | MedChem Express | HY-N0504 | 75330-75-5 |
| BMS-303141 | MedChem Express | HY-16107 | 943962-47-8 |
| CTPI-2 | MedChem Express | HY-123986 | 68003-38-3 |
| Ferrostatin-1 | MedChem Express | HY-100579 | 347174-05-4 |
| Sodium citrate | Sigma | PHR1416-1G | - |

Table S4:antibodies used for Flow Cytometry

| Antibody | Fluorophore | Supplier | Cat# |
| --- | --- | --- | --- |
| Rabbit monoclonal anti-H3K27me3 | PE-Cy7 | Cell Signaling Technologies | 91611S |
| Rabbit polyclonal anti-H4K16ac | / | Active Motif | 39068 |
| Rabbit polyclonal anti-H3K9ac | / | Active Motif | 39038 |
| Rabbit anti-Histone 3 | / | Cell Signaling Technologies | 9715S |
| Rabbit anti-SQLE | / | Cell Signaling Technologies | 40659S |
| Rabbit anti-SAT1 | / | Cell Signaling Technologies | 61586S |
| Rabbit anti-H3K27ac [Clone D5E4] | Alexa Fluor 647 | Cell Signaling Technologies | 39030S |
| Rabbit anti-MYB | / | Cell Signaling Technologies | 12319S |
| Rabbit anti-MYB | PE | Cell Signaling Technologies | 73288S |
| Goat anti-Rabbit IgG | Alexa Fluor 405 | Abcam | ab175652 |
| Goat anti-Rabbit IgG | Alexa Fluor 467 | Abcam | ab150079 |

Table S5: antibodies used for Western Blotting

| Antibody | Supplier | Cat# |
| --- | --- | --- |
| Mouse monoclonal Anti-β–Actin [Clone AC-15] | Merck | A5441 |
| Rabbit anti-H3K27ac | Cell Signaling Technologies | 8173S |
| Rabbit anti-MYB [Clone D2R4Y] | Cell Signaling Technologies | 12319S |
| Rabbit anti-HMGCS1 [Clone D1Q9D] | Cell Signaling Technologies | 42201S |
| Rabbit anti-SQLE | Cell Signaling Technologies | 40659S |
| Rabbit anti-GPX4 | Cell Signaling Technologies | 52455S |

Table S6: primers used for qRT-PCR

| Target | Primer direction | Sequence |
| --- | --- | --- |
| EZH2 | Fw | TATTCTTGGTCTCCCCTACAGC |
|  | Rv | TCTTCAATGAAAGTACCATCCTG |
| ACLY | Fw | AGACCTCGATGCCAAAAGTG |
|  | Rv | TAGTTTGCCAGCTCGTTGAC |
| FASN | Fw | ACCTCCGTGCAGTTCTTGAG |
|  | Rv | AGGGACTTCTTGGTCAGCAG |
| SQLE | Fw | GTGATGGGAGTTCAGTACAAGG |
|  | Rv | ATTGGAGACCAGGCTTTTCC |
| HMGCS1 | Fw | CTCGGATGTTGCTGAATGAC |
|  | Rv | GCCTTCATAAATGCCTTCTCC |
| HMGCR | Fw | TTCTTCCCAGCTTGTGTGTC |
|  | Rv | ACCCTCTGAGTTACAGGATTCG |
| SAT1 | Fw | TTACCTATGACCCGTGGATTG |
|  | Rv | TTTCTGATCCTATGCCAAAGC |
| BCL2 | Fw | CTGAGTACCTGAACCGGCA |
|  | Rv | CTGAGTACCTGAACCGGCA |
| BCL2L1 | Fw | CCTAAGGCGGATTTGAATCTCT |
|  | Rv | TGGGCTCAACCAGTCCATTG |
| MYB | Fw | CCGTTTTAATGGCACCAGCA |
|  | Rv | CCCAGGTACTGCTACAAGGC |
